# Reduced levels of synaptic vesicle protein 2A in the extracellular vesicles and brain of Alzheimer’s disease- associations with Aβ, tau and synaptophysin

**DOI:** 10.1101/2025.01.21.634009

**Authors:** Jana Nussbaumer, Aatmika Barve, Valentin Zufferey, Jeanne Espourteille, Tunahan Kirabali, Uwe Konietzko, Daniel Razansky, Axel Rominger, Agneta Nordberg, Luc Buée, Morvane Colin, Roger M. Nitsch, Christoph Hock, Kevin Richetin, Ruiqing Ni

## Abstract

**Background:** Synaptic dysfunction plays an important role in Alzheimer’s disease (AD) and is an emerging imaging and fluid biomarker. Here, we aimed to assess the regional expression of synaptic vesicle glycoprotein 2A (SV2A) in the brain and extracellular vesicles of AD patients and its associations with the APOE ε4 allele, amyloid-β, tau pathologies, and other synaptic markers.

**Methods:** Mass spectrometry-based synaptosome proteomics was performed on brain-derived extracellular vesicles (BdEVs) isolated from the frontal cortex of 17 AD patients and 4 NCs. Immunohistochemical staining for SV2A, synaptophysin, amyloid-β and phospho-tau was performed on postmortem tissue from the frontal, temporal, and entorhinal cortices and hippocampus of 40 AD patients and 44 nondemented controls (NCs).

**Results:** Reduced levels of synaptic proteins, including synaptotagamin, GAP43, SYT1, SNAP25 and 14-3-3ζ, were positively correlated with SV2A and negatively correlated with GFAP and NEFL in BdEVs from AD patients and NCs. We detected lower levels of SV2A in the hippocampus and entorhinal cortex of AD compard to NCs, and in APOE ε4 carriers than in noncarriers. SV2A levels were positively correlated with synaptophysin and negatively correlated with the levels of the amyloid-β, phospho-tau, and Braak stages.

**Conclusions:** This study provides postmortem evidence of synaptic markers and reduced regional levels of SV2A in brain tissue slices and BdEVs from AD patients compared with NCs and in APOE ε4 carriers compared to non-carriers. SV2A could serve as a valuable marker for monitoring synaptic degeneration in AD.

## Introduction

Alzheimer’s disease (AD) is a neurodegenerative disorder characterized by progressive memory loss and cognitive decline. Clinically, the disease can be divided into three stages: preclinical AD or cognitive unimpaired people with amyloid-β accumulation in the brain; prodromal or mild cognitive impairment (MCI); and AD dementia (25). AD is pathologically characterized by the abnormal accumulation of amyloid-β (Aβ) plaques and neurofibrillary tangles (NFTs) formed by hyperphosphorylated tau (6, 28). Synapse damage and loss are downstream effects of amyloidosis and tauopathy, lead to neurodegeneration and are recognized as one of the earliest neurobiological correlates of cognitive impairment in AD (11, 30, 40, 57, 71, 77). Aβ oligomers, rather than plaques, have been shown to induce synapse degeneration in AD animal models (61). Neuropathological studies have shown that synaptic protein expression is lower in NFT-containing neurons than in those without NFT, which is correlated with the Braak stage and the level of NFT (15, 23, 59). Pathogenic tau binds to synaptic vesicles, dysregulates the synaptic proteome and interferes with presynaptic functions, synaptic transmission, synaptic vesicle mobility and the release rate, leading to synaptic loss (58, 73, 84, 87). As pathological tau spreads transsynaptically in a ‘prion-like’ manner, the deposition of hyperphosphorylated tau at multiple points can consequently spread along axons and lead to pronounced synaptic loss in widespread regions. An earlier study demonstrated that synaptic oligomeric tau is a likely culprit in the spread of tau pathology through the brain in AD patients via subdiffraction-limited microscopy (10). In addition, microglia and astrocytes drive synaptic degeneration in animal models of AD via the activation of complement receptors and elimination of synaptic structures, leading to synapse loss, neuronal dysfunction and cognitive decline (17, 22, 68).

Synaptic biomarkers might be used in biomarker panels for diagnostics, disease staging, and prediction of progression and have shown potential as tools to monitor downstream effects on synaptic function and the integrity of treatments in drug trials (48, 65). Positron emission tomography (PET) using synaptic vesicle glycoprotein 2A (SV2A) radioligands ([^11^C]UCB-J, [^18^F]UCB-H, and [^18^F]SynVesT-1, [^18^F]SynVesT-2) (8, 18, 29, 43, 49) has been developed for in vivo imaging. SV2A is expressed at presynaptic terminals throughout the brain, can influence neurotransmitter release by regulating the amount of synaptotagmin (SYT1) in secretory vesicles, can be important in the trafficking of SYT1 to synaptic vesicles and can regulate the effectiveness of calcium in inducing vesicle fusion. Synaptophysin (SYP) is an integral synaptic vesicle protein that modulates endocytosis and is crucial for trafficking the essential vesicular SNARE protein synaptobrevin II (33, 75). A reduction in SV2A has been demonstrated in the hippocampus and cortical regions of amnesic mild cognitive impairment (aMCI) patients with AD compared with nondemented controls (NCs) and is related to tau deposition (74). In addition, hippocampal SV2A tracer uptake and serum levels of SV2A are positively correlated with cognitive performance in patients with AD and decrease with the progression of AD (3, 11, 44, 80). In addition, fluid biomarkers (CSF and blood) based on ultrasensitive immunoassays, such as synaptosomal-associated protein 25 kDa (SNAP-25), neurogranin, visinin-like protein 1 (VILIP-1), neuronal pentraxin 2, β-synuclein, SYT1, and growth-associated protein 43 (GAP43), have been identified (19, 38, 85). Mass spectrometry-based proteomic analysis of CSF and plasma has demonstrated differentiation between sporadic AD patients and NCs as well as between sporadic AD patients and autosomal dominant AD patients (62, 67). In addition, emerging evidence has shown the potential of blood extracellular vesicles (EVs) carrying synaptic function- and brain-related proteins as potential biomarkers for AD (70).

The aims of this study were to 1) map alterations in the levels of SV2A, SYP and other synaptic proteins in the brains of AD patients compared with NCs; 2) assess the associations between SV2A or SYP and the Aβ, tau, and APOE ε4 alleles in the brains of AD patients and NCs; and 3) understand the associations between SV2A and other synaptic markers in the EV. We included 105 cases and performed mass spectrometry-based synaptosome proteomics on brain-derived EVs (BdEVs) and immunohistochemical and immunofluorescence staining of postmortem brain tissue.

## Materials and methods

### Postmortem human brain tissue

Autopsy frozen frontal cortex tissue from 17 AD patients and 4 NC patients for BdEV extraction and proteomics was obtained from CHUV, Switzerland, as described previously (36, 56). Postmortem paraffin-embedded brain tissue from 40 patients with AD, each with a clinical diagnosis confirmed by pathological examination, and 44 NCs were obtained from the Netherlands Brain Bank (NBB), Netherlands, for IHC/IF studies (**Table 1**, including the hippocampus, EC, frontal cortex and temporal cortex). All materials were collected from donors or from whom written informed consent was obtained for a brain autopsy, and the use of the materials and clinical information for research purposes were obtained by the NBB or CHUV. The study was conducted according to the principles of the Declaration of Helsinki and subsequent revisions. All the autopsied human brain tissue experiments were carried out in accordance with ethical permission obtained from the regional human ethics committee and the medical ethics committee of the VU Medical Center for NBB tissue. Information on APOE ε4 and the neuropathological diagnosis of AD (possible, probable, or definite AD) or not AD was obtained from the NBB. Information on the Consortium to Establish a Registry for AD (CERAD), which applies semiquantitative estimates of neuritic plaque density and the Braak stage (6) on the basis of the presence of neurofibrillary tangles (NFTs), was obtained from the NBB and CHUV and is provided in **Table 1**.

**Table 1.**
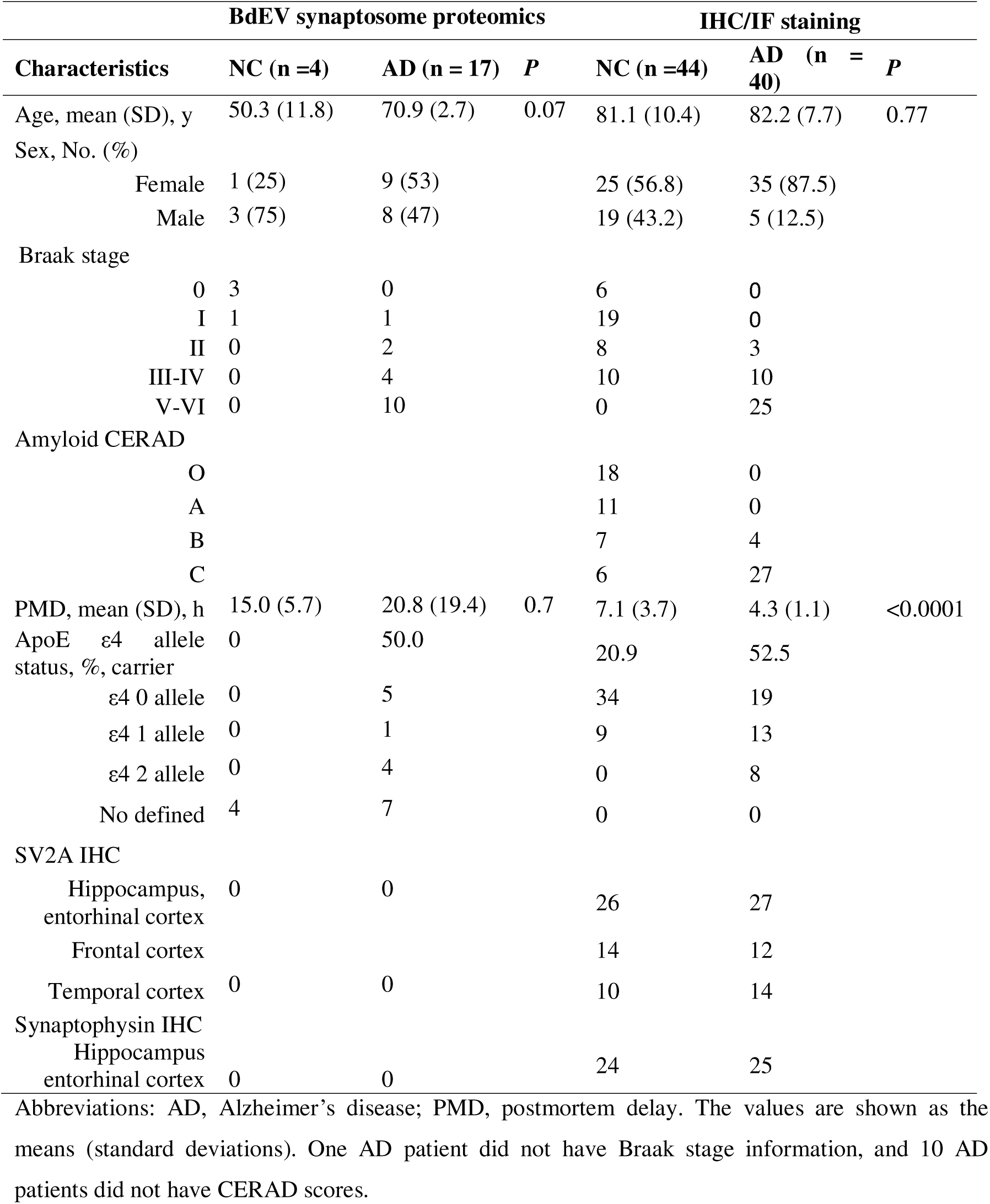
Demographic and neuropathologic characteristics of postmortem AD patients and NCs in proteomics and in staining.

### Interstitial fluid and BdEV isolation from patient prefrontal cortex tissue

Brain-derived fluids (Bd-fluids) were isolated as previously described (36, 56). Briefly, approximately 1.5 g of fresh-frozen patient prefrontal cortex was gently cut into smaller pieces. These samples were then incubated in a Papain/Hibernate-E solution to gently digest the tissue and release interstitial fluid, theoretically avoiding cellular lysis. After tissue digestion, differential centrifugation at 300 × g for 10 min, 2,000 × g for 10 min, and 10,000 × g for 30 min was performed at 4°C to remove cells, membranes, and debris, respectively, and the supernatant from the 10,000 × g centrifugation was then stored at -80°C until EV isolation was performed. To avoid bias in the results, normalization according to the weight of the brain extracts was systematically performed. The procedures used to isolate BdEVs from human BD fluid were carried out in accordance with the minimal information for the studies of extracellular vesicles (MISEV) guidelines that were established and updated in 2024 by the International Society for Extracellular Vesicles (82). Various controls were used to validate the enrichment and content of the BdEVs, as recommended in these guidelines. However, the procedure described above to recover BD fluids may still lead to some cell lysis and therefore some intracellular contamination. The presence of intraluminal vesicles (ILVs) in the preparations cannot be fully excluded. A total of 500 μL of BD fluid was loaded on top of a size exclusion chromatography (SEC) column (10 mL column, CL2B Sepharose, pore size 75 nm, Millipore). A mean of 7.94 × 10^10^ (±3.36 × 109 vesicles/g of tissue in F1–4 (n = 36 samples) was recovered for each sample. Isolation was carried out in phosphate-buffered saline (PBS) with a flow of 36–48 s/mL. The first 3 mL was removed, and the following 20 fractions were recovered (with 500 μL per fraction).

### Label-free liquid chromatography (LC) tandem mass spectrometry (MS)

F1l4 fractions were digested according to a modified version of the iST method (named the miST method) (31, 36). Briefly, 50 mL of solution in PBS was supplemented with 50 mL of miST lysis buffer (1% sodium deoxycholate, 100 mM Tris pH 8.6, 10 mM dithiothreitol) and heated at 95°C for 5 min. Samples were then diluted 1:1 (v:v) with water, and reduced disulfides were alkylated by adding 1/4 vol of 160 mM chloroacetamide (final 32 mM) and incubating at 25°C for 45 min in the dark. The samples were adjusted to 3 mM ethylenediaminetetraacetic acid and digested with 0.5 mg of trypsin/LysC mixture (Promega #V5073) for 1 h at 37°C, followed by a second 1 h of digestion with a second and identical aliquot of proteases. To remove sodium deoxycholate and desalted peptides, two sample volumes of isopropanol containing 1% TFA were added to the digests, and the samples were desalted on a strong cation exchange (SCX) plate (Oasis MCX; Waters Corp., Milford, MA) by centrifugation. After washing with isopropanol/1% trifluoroacetic acid, the peptides were eluted in 250 mL of 80% acetonitrile, 19% water, and 1% (v/v) ammonia. Tryptic peptide mixtures were then injected on an Ultimate RSLC 3000 nanoHPLC system interfaced via a nanospray Flex source to a high-resolution Orbitrap Exploris 480 mass spectrometer (Thermo Fisher, Bremen, Germany). Peptides were loaded onto an Acclaim PepMap100 C18 microcolumn (20 mm × 100 μm ID, 5 μm, Dionex) before separation on a C18 custom-packed column (75 μm ID × 45 cm, 1.8 μm particles, Reprosil Pur, Dr. Maisch), and a gradient from 4 to 90% acetonitrile in 0.1% formic acid was used for peptide separation (total time: 140 min). Full MS survey scans were performed at 120,000 resolution. A data-dependent acquisition method controlled by Xcalibur software (Thermo Fisher Scientific) was used to optimize the number of precursors selected ("top speed") to charge 2+ to 5+ while maintaining a fixed scan cycle of 2 s. Peptides were fragmented by higher energy collision dissociation with a normalized energy of 30% at 15000 resolution. The window for precursor isolation was 1.6 m/z units around the precursor, and selected fragments were excluded for 60 s from further analysis.

### MS and MS data analysis

The data files were analyzed with MaxQuant 1.6.14.0, which incorporates the Andromeda search engine (13). Cysteine carbamidomethylation was selected as a fixed modification, whereas methionine oxidation and protein N-terminal acetylation were specified as variable modifications. The sequence databases used for searching were the human (*Homo sapiens*) reference proteome from the UniProt database and a "contaminant" database containing the most common environmental contaminants and enzymes used for digestion (keratins, trypsin, etc.). The mass tolerance was 4.5 ppm for the precursors (after recalibration) and 20 ppm for the MS/MS fragments. Both peptide and protein identifications were filtered at a 1% false discovery rate relative to hits against a decoy database built by reversing protein sequences. Proteins were annotated according to several databases. These databases include the Gene Ontology (GO) Human database, which is further split into the categories of cellular component (CC), biological process (BP), and molecular functions (MF), as well as the MISEV guidelines for EV-associated and potential contaminant proteins, a database generated by McKenzie and colleagues in their publication for brain cell type specificity, and the Human Protein Atlas databases for cell type specificity and predicted subcellular localization. Within the neuronal category, we also compared excitatory and inhibitory neuron-specific proteins according to the Human Protein Atlas (1).

### Weighted gene coexpression network analysis

Protein coexpression network analysis was performed with the R package weighted gene coexpression network analysis (WGCNA) (35) on the preprocessed proteomic data of all identified proteins with their respective riBAQ scores for each disease group. First, a correlation matrix for all pairwise correlations of proteins across all samples was generated and then transformed into a matrix of connection strengths, i.e., a weighted adjacency matrix with soft threshold power β = 16. The topological overlap (TO) was subsequently calculated via the connection strengths. Proteins were further hierarchically clustered via the 1-TO measure as the distance measure to generate a cluster dendrogram, and modules of proteins with similar coexpression relationships were identified via the dynamic tree-cutting algorithm with the following parameters: minimal module size = 30, deepSplit = 2, and merge cut height = 0.15. Within each module, the module eigenprotein was defined as the first principal component, which serves as a weighted summary of protein expression within the module and accounts for the greatest variance among all module proteins. Furthermore, module membership (kME) was assigned by calculating Pearson’s correlation between each protein and each module eigenprotein and the corresponding p value. Proteins were reassigned to the module for which they had the highest module membership, with a reassignment threshold of p < 0.05. From each module, hub proteins were identified via the signedkME function, which explains the membership of a protein with its module and its strong association within the module.

### Gene ontology

A detailed genetic annotation of each protein within the WGCNA modules was performed. These differentially expressed proteins and coexpressed proteins were characterized on the basis of their gene ontologies via the GO-Elite (version 1.2.5) python package (86). The entire set of proteins identified and included in the network analysis served as the background dataset. The presence of significantly overrepresented ontologies within a module was gauged via a Z score, whereas the significance of the Z scores was evaluated via a one-tailed Fisher’s exact test, with adjustments for multiple comparisons via the BenjaminilHochberg false discovery rate method. The threshold analysis included a Z score cutoff of 2, a p value threshold of 0.05, and a minimum requirement of five genes per ontology before ontology pruning was performed. The best gene ontology term, which explains the molecular and cellular functions of each module, was used for naming.

### Protein**□**protein interaction network

Proteins used as inputs for network generation were selected from significant WGCNA modules. We used the SIGNOR database (ref SIGNOR app + specific tool used) to generate an interaction network in Cytoscape (v.3.10) for each significant module (39). The obtained networks were merged into a single network that was currated to keep only proteins present in our data and the most prominent subnetworks. Finally, information on module appartenance and correlation with SV2A was added to improve the visual representation and relevance of the network.

### Immunohistochemistry, immunofluorescence staining, microscopy and image analysis

The paraffin-embedded fixed postmortem brain tissues (from 40 AD patients and 44 NCs) were cut into 3 µm sections via a Leica microtome (Leica Microsystems, Germany). Hematoxylin and eosin (H&E) staining was performed according to routine procedures for each patient to provide anatomical information and to determine whether there were abnormalities in the brain. Immunochemical staining with antibodies against SV2A, SYP, and Aβ_17-24_ (4G8) was performed in the hippocampus, entorhinal cortex, frontal cortex and temporal cortex (27, 52). Immunofluorescence staining using antibodies against phospho-Tau (AT8, Ser202/Thr205) was performed in the hippocampus and entorhinal cortex. To assess the spatial associations between SV2A and Aβ and tau, double immunofluorescence staining was performed using SV2A/6E10 (antibodies against Aβ_1-16_) and SV2A/AT8 on frontal cortex, temporal cortex and hippocampus/entorhinal cortex slices from AD patients and NCs. The details of the antibodies, kits and chemicals used are listed in **STable 1**. Paraffin-embedded fixed human brain tissue sections were permeabilized and blocked in 5% normal donkey or goat serum and 1% Triton-phosphor-buffered saline (PBS) for one hour at room temperature with mild shaking. The slices were then incubated with primary antibodies overnight at 4°C with mild shaking. The next day, the sections were washed with PBS two times for 20 minutes and incubated with a suitable secondary antibody for 2 hours at room temperature. The sections were incubated for 15 minutes in 4′,6-diamidino-2-phenylindole (DAPI), washed two times for 10 minutes with 1×PBS, and mounted with VECTASHIELD Vibrance Antifade Mounting Media (Vector Laboratories, Z J0215). The brain sections were imaged at ×20 magnification via an Axio Oberver Z1 slide scanner (Zeiss, Germany) and ×20 magnification via Leica SP8 confocal microscopy (Leica, Germany) via the same acquisition settings.

For the analysis of the staining results, manual delineation of regions of interest (ROIs) for the hippocampus CA1, CA2/3, dentate gyrus (DG), subiculum (SUB), entorhinal cortex (EC), gray matter and white matter in the frontal and temporal cortex was performed using the Allen atlas as a reference (16). Qupath and ImageJ (NIH, U.S.A.) were used for image processing and analysis. The percentage of positively stained area was quantified for immunochemical staining with SV2A, SYP, and 4G8 antibodies, and the mean fluorescence intensity was quantified for AT8 staining (41).

### Statistics

All the statistical analyses were performed via GraphPad Prism 10 (GraphPad). The ShapirolWilk test was used to assess the normality of the distributions of various parameters in both groups. For the statistical analysis of differences in the numerical variables between two groups, the nonparametric MannlWhitney U test was used. The chi-square test was used to assess differences in categorical variables between two groups. Nonparametric Spearman’s rank correlation analysis was performed for those that failed the normality test. Significance was set at p <0.05. The values are shown as the mean ± standard deviation.

## Results

### Demographics and pathological description

The details of the characteristics of the patients included are presented in **Table 1**. The ages of the individuals in the NC group (81.1±10.4, n=44) were comparable to those in the AD group (82.2±7.7, n=40) and followed a normal statistical distribution (ShapirolWilk test). A greater percentage of females was noted in the AD group (85.5%) than in the NC group (56.8%). The prevalence of APOE ε4 carriers was greater in the AD group (52.5%) than in the NC group (20.9%). There were no APOE 2ε4 carriers in the NC group. H&E staining revealed no abnormalities in the brain tissue slices from the AD patients or NCs (examples are shown in **SFig. 7**). The postmortem delay of brain tissues was rather short for the NC (7.1±3.7 hours) and AD (4.3±1.1 hours) groups, which ensures suitability for synaptic protein analysis. The tissues used for proteomics analysis are shown in Table 1. The ages of the individuals in the NC group (50.3±11.8, n=4) were not significantly different from those in the AD group (70.9 ±2.7, n=17).

We used 4G8 immunohistochemical staining to map the Aβ distribution in the hippocampus, entorhinal cortex, frontal cortex and temporal cortex slices and AT8 immunofluorescence staining to map the pathological phospho-Tau distribution in the hippocampus and entorhinal cortex slices (**Fig. 6, SFigs. 8**). Aβ plaques and tau pathology were also observed in the brains of a proportion of NCs (**Table 1**).

### Compared with that in NCs, SV2A expression in BdEVs was positively correlated with other synaptic proteins in AD and NC brains

Following a previously published protocol (36, 56), we isolated BdEVs from brain biofluids collected from the prefrontal cortex of 4 NCs and 17 patients with AD at different stages. We employed a previously published method, combining slow digestion of freshly frozen patient tissues with size-exclusion chromatography (36). A portion of the isolated vesicles was analyzed via nanoparticle tracking analysis to quantify their concentration and size, whereas another fraction was digested with trypsin for in-depth analysis via LC-MS/MS (**Fig. 1a**). Our results revealed that neither the number of vesicles (**Fig. 1b**) nor the number of proteins detected (**Fig. 1c**) varied significantly between the NC and AD groups, regardless of disease stage. Annotation of the identified proteins according to the MISEV2024 guidelines further confirmed that the relative abundance of extracellular vesicle-associated proteins, as well as potential contaminants, remained stable across groups (**SFig. 1)**. These initial quality control analyses indicate that the progression of AD does not impact the quality of the BdEVs isolated from the prefrontal cortex of AD patients. To investigate the global proteomic signature of BdEVs, we applied a correlation-based approach involving WGCNA, which identified eight functional modules (**Fig. 1d-l**). Among these modules, the M2 ("Cell signaling") (**Fig. 1f**) and M3 ("Synaptic vesicles") (**Fig. 1g**) modules were significantly reduced in AD patients. In contrast, the M6 ("endoplasmic reticulum") (**Fig. 1i**) and M7 ("axonal transport") (**Fig. 1k**) modules significantly increased at advanced disease stages (Braak V-VI). The M3 module was the only module significantly altered from the early stages of the disease and across all stages, suggesting a strong association between BdEV proteomic signatures and AD progression, particularly concerning synaptic proteins.

**Figure 1:**
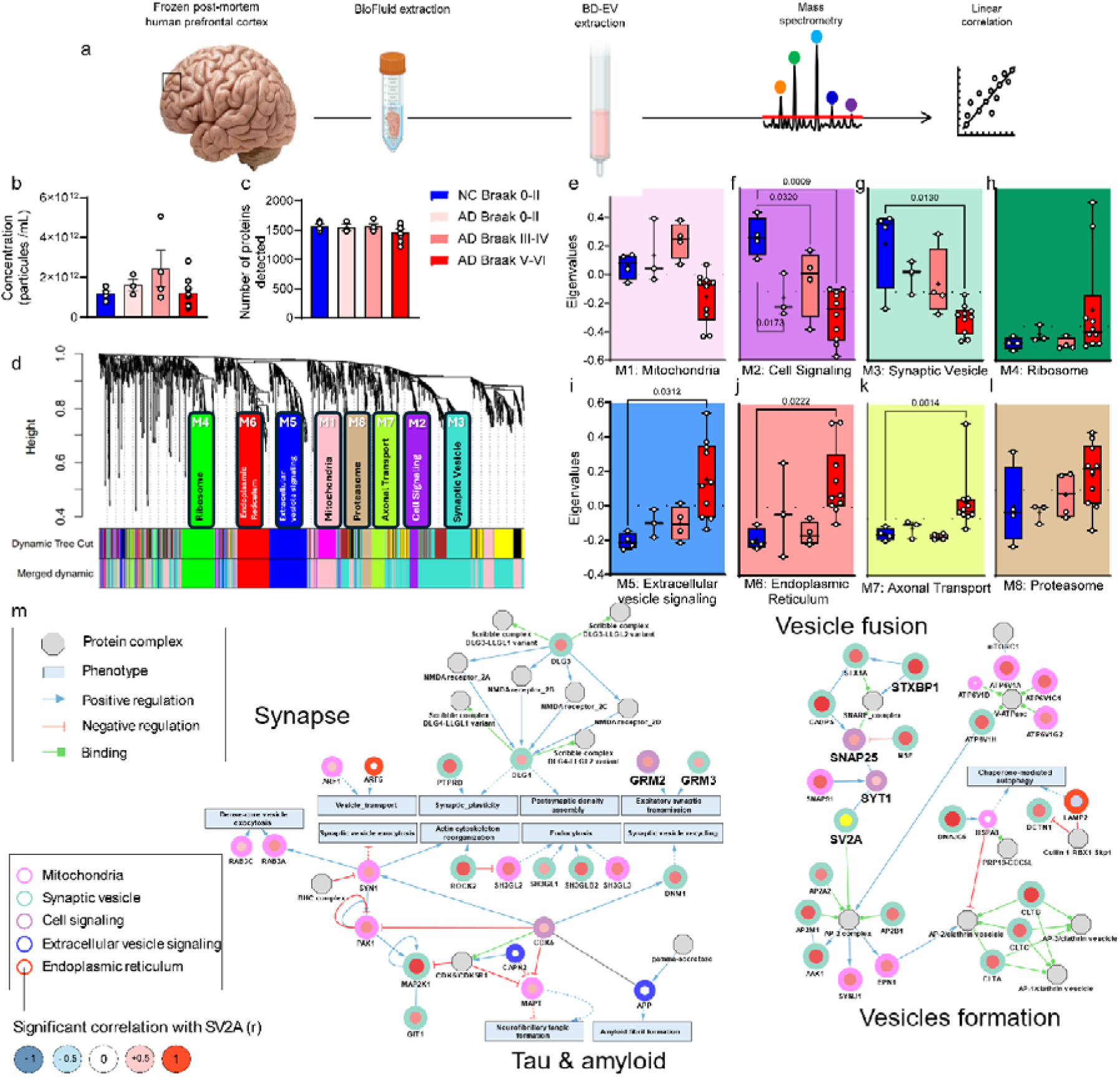
Proteomics of BdEV from AD and NC brains. (a) Scheme of the experimental design for brain-derived EV proteomics. (b) Histogram showing the relative quantities (rIBAQs) of concentration particles between the NCs and ADs of different braak stages. (c) Histogram showing the relative quantities (rIBAQs) of protein between the NCs and ADs of different Braak stages. (d) WGCNA cluster dendrogram and significant modules. (e-l) Comparisons of module eigenvalues between NCs and Ads of different Braak stages for mitochondria (e), cell signaling (f), synaptic vesicles (g), ribosomes (h), extracellular vesicle signaling (i), the endoplasmic reticulum (j), axonal transport (k), and proteasomes (l). (m) Proteinlprotein interaction network for proteins of the WGCNA modules (circles) with additional protein complexes (octagons) and phenotypes (rectangles). Border color indicates the module of the protein, and fill color indicates the r of correlative analysis (Pearson’s correlation; red=positive r>0.5; blue=negative, r<0.5). The color of the edge indicates the type of interaction (black: uinspecified, Blue, blue: positive regulation, red: negative regulation, and green: binding). The dashed lines indicate indirect interactions.

To further explore the modules, we analyzed protein interactions via the Signaling Network Open Resource (SIGNOR) platform (**Fig. 1m**). Two main subnetworks were identified, highlighting proteins involved in synaptic functions, vesicular transport, and proteopathic mechanisms related to tau and Aβ. Within the M3 module, the protein SV2A emerged as a central hub owing to its numerous interactions with proteins from subnetworks related to vesicle formation and fusion. To better understand the role of SV2A in BdEVs, we conducted linear correlation analyses between its relative abundance and that of other proteins in the modules. We found that many proteins within associated clusters were positively correlated with SV2A (red center nodes). These proteins were predominantly located within the "Synaptic vesicles", "Mitochondria", and "Cell signaling" modules (turquoise, pink, and purple borders, respectively). Finally, to assess whether SV2A and other synaptic markers are reduced in BdEVs and whether SV2A levels are correlated with other recently proposed CSF panel biomarkers of synaptic dysfunction in neurodegenerative diseases (53), we performed nonparametric Spearman rank correlation analyses. Among this panel of 24 proteins, 21 were detectable in BdEVs (**Fig. 2, SFig. 2, Table 2**). Notably, we observed significant decreases in proteins at the late stage of AD (AD Braak stages V-VI), including adaptor-related protein complex 2 subunit beta 1 (AP2B1), complex 2 (CPLX2), GAP43, SNAP25, gamma-synuclein (SNCG), SV2A, syntaxin 1B (STX1B), SYP; synaptotagmin 1 (SYT1), tubulin teta 3 class III (TUBB3) and 14-3-3ζ (YWHAZ) (**Fig. 2a-l**). Interestingly, only GAP43 and SYT1 were significantly reduced in BdEVs at the early stages of the disease (AD Braak stages III-IV) (**Fig. 2c, j**). In contrast, no differences in the levels of GDP dissociation inhibitor 1 (GDI1), glial fibrillary acidic protein (GFAP), neuronal pentraxin 1 (NPTX1), neurofilament light chain (NEFL), phosphatidylethanolamine binding protein 1 (PEBP1), STY7 or postsynaptic metabotropic glutamate receptor subtype 5 (GRM5) were detected between AD patients and NCs (**SFig. 2**)(78).

**Fig. 2.**
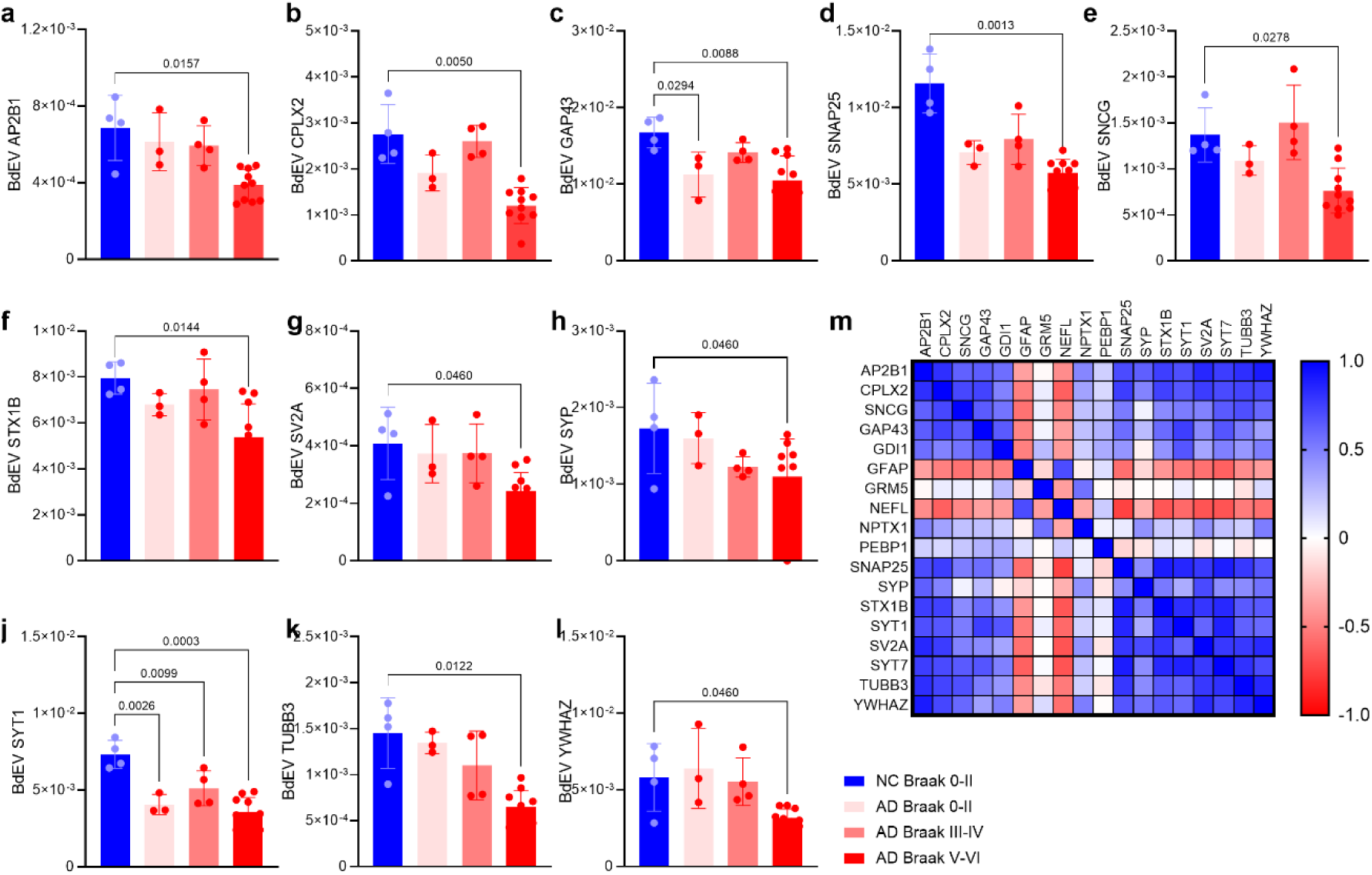
Comparison and correlation between cortical synaptosome BdEV markers in NC and AD. (a-l) Comparison of BdEV levels of AP2B1, CPLX2, GAP43, SNAP25, SNCG, STX1B, SV2A, SYP, SYT1, TUBB3 and YWHAZ in the NC Braak 0-II, AD of Braak 0-II, III-IV, and V-VI. (m) Nonparametric Spearman rank analysis of the rIBAQ matrix of correlation in the AD and NC groups (detailed p values and r values are provided in Table 2, and supplemental files).

**Table 2.** Nnparametric Spearman rank correlation analysis between SV2A, SYP and other markers in the BdEV of AD and NC brains.

| Gene Name | Detectable | SV2A |  |  |  | SYP |  |  |  |
| --- | --- | --- | --- | --- | --- | --- | --- | --- | --- |
|  |  | AD+NC |  | AD |  | AD+NC |  | AD |  |
|  |  | R | P value | R | P value | R | P value | R | P value |
| AP2B1 | Yes | 0.8460 | **<0.0001** | 0.8460 | **<0.0001** | 0.6273 | **0.0023** | NS |  |
| CHAT | No |  |  |  |  |  |  |  |  |
| CHRNA7 | No |  |  |  |  |  |  |  |  |
| CMPLX2 | Yes | 0.7609 | **0.0001** | 0.7609 | **0.0001** | 0.4857 | **0.0256** | NS |  |
| GAP43 | Yes | 0.4587 | **0.0365** | 0.4587 | **0.0365** | NS |  | NS |  |
| GDI-1 | Yes | 0.3502 | 0.1196 | 0.3502 | 0.1196 | NS |  | NS |  |
| GFAP | Yes | -0.5608 | **0.0082** | -0.5608 | **0.0082** | NS |  | NS |  |
| GRM5 | Yes | -0.0023 | 0.9921 | -0.0023 | 0.9921 | NS |  | NS |  |
| NEFL | Yes | -0.6725 | **0.0008** | -0.6725 | **0.0008** | NS |  | NS |  |
| NPTX1 | Yes | 0.4228 | 0.0562 | 0.4228 | 0.0562 | NS |  | NS |  |
| NPTX2 | Yes |  | NS |  | NS | NS |  | NS |  |
| NPTXR | Yes |  | NS |  | NS | NS |  | NS |  |
| NRGN | No |  |  |  |  |  |  |  |  |
| PEBP-1 | Yes | -0.0981 | 0.6722 | -0.0981 | 0.6722 | NS |  | NS |  |
| SNAP-25 | Yes | 0.7843 | **<0.0001** | 0.7843 | **<0.0001** | 0.4494 | **0.0409** | NS |  |
| SNCB | No |  |  |  |  |  |  |  |  |
| SNCG | Yes | 0.4873 | **0.0250** | 0.4873 | **0.0250** | NS |  | NS |  |
| STX1B | Yes | 0.7888 | **<0.0001** | 0.7888 | **<0.0001** | 0.5247 | **0.0146** | NS |  |
| SV2A | Yes | / | / | / | / | 0.7063 | **0.0003** | 0.5236 | **0.0328** |
| SYP | Yes | 0.7063 | **0.0003** | 0.5236 | **0.0328** | NS |  | NS |  |
| SYT-1 | Yes | 0.6023 | **0.0039** | 0.6023 | **0.0039** | NS |  | NS |  |
| SYT-7 | Yes | 0.7781 | **<0.0001** | 0.7781 | **<0.0001** | 0.4434 | **0.0441** | NS |  |
| TUBB3 | Yes | 0.7797 | **<0.0001** | 0.7797 | **<0.0001** | 0.4532 | **0.0391** | NS |  |
| VILIP-1 | No |  |  |  |  |  |  |  |  |
| 14-3-3ζ | Yes | 0.8369 | **<0.0001** | 0.8369 | **<0.0001** | 0.5519 | **0.0095** | NS |  |

AP2B1, adaptor-related protein complex 2 subunit beta 1; CPLX2, Complexin 2; GAP43, growth-associated protein 43; SNAP25, synaptosomal-associated protein 25 kDa; SNCG, gamma-synuclein; SV2A, synaptic vesicle glycoprotein 2A; STX1B, syntaxin 1B; SYP; SYT1 or 7, synaptotagmin 1 or 7; TUBB3, tubulin teta 3 class III; YWHAZ, 14-3-3ζ; GDI1, GDP dissociation inhibitor 1; GFAP, glial fibrillary acidic protein; NPTX1, neuronal pentraxin 1; NEFL, neurofilament light chain; PEBP1, phosphatidylethanolamine binding protein 1; GRM5, metabotropic glutamate receptor subtype 5;

The different BdEV markers analyzed were correlated with the AD+NC group (**SFig. 3**) and the AD group (**SFig. 4**). Among these 24 proteins, 13 were positively correlated with the relative levels of SV2A, with AP2B1, YWHAZ, SNAP25, STX1B, STY7, TUBB3 and CPLX2 showing the highest correlations (R>0.75, p<0.0001) (**Fig. 2**, **Table 2, SFig. 5**), indicating synaptic and neuronal loss and concomitant astrogliosis. In comparison, the relative levels of SYP were correlated with those of 7 proteins (all R values<0.75). These data suggest that the sysnaptic markers in BdEV are particularly central in detecting the progression of the pathology and that the SV2A protein seems to be central and correlates well with the other proteins, suggesting that SV2A could serve as a valuable marker for monitoring synaptic degeneration in AD.

### Reduced levels of SV2A in AD patients compared with NCs and in APOE **ε**4 allele carriers compared with noncarriers

We first quantified SV2A expression in brain tissue slices from the hippocampus, entorhinal cortex, frontal cortex, and temporal cortex of AD patients and NCs via immunochemical staining. The SV2A levels did not differ significantly among subfield regions of the hippocampus. Compared with those in NCs, SV2A levels were lower in all the hippocampal subfields of AD patients, namely, the DG (p=0.0017), CA2/3 (p=0.0462), CA1 (p=0.0024), and Sub (p=0.0462), as well as in the entorhinal cortex (p=0.0462) (**Fig. 3a-f**, **Fig. 4a, SFig. 6**). In the frontal cortex and temporal cortex of both AD patients and HCs, the level of SV2A in gray matter was several-fold greater than that in white matter (**Figs. 3g-j**, **Fig. 4d, e**). The levels of SV2A in the gray matter or white matter in the frontal cortex and temporal cortex of AD patients were comparable to those in NC patients on the basis of immunohistochemical staining (**Fig. 3, 4d, e, SFig. 6**). The laminar distribution of SV2A in the gray matter of the frontal, temporal and entorhinal cortex is rather homologous. In contrast, the pyramidal layer of the hippocampus presented a dense SV2A signal.

**Fig. 3.**
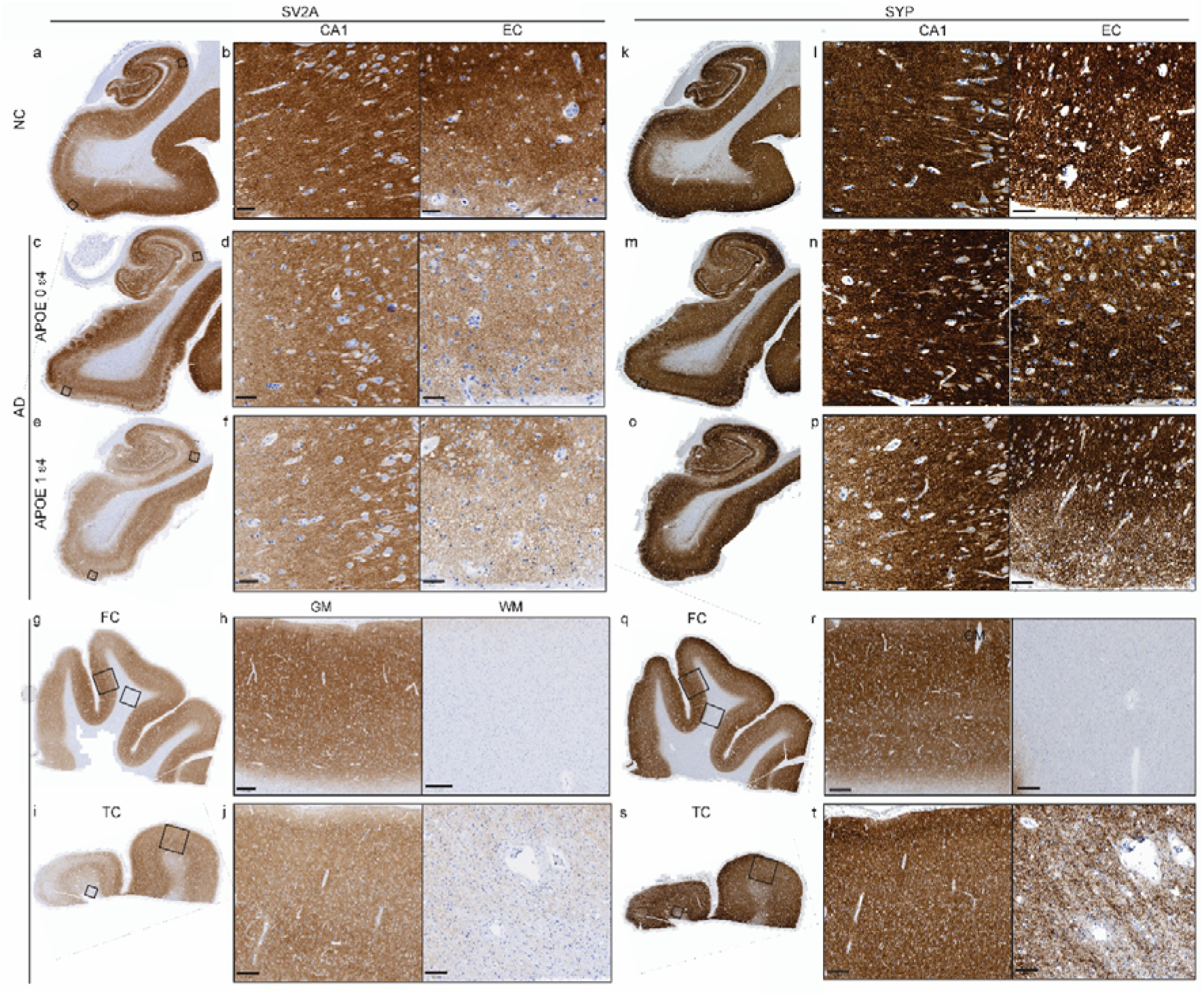
SV2A and SYP staining in the hippocampus, entorhinal cortex, frontal cortex and temporal cortex of NC and AD APOE ε4 carriers and noncarriers. (a-f, k-p) Representative images of immunohistochemical staining for SV2A (a-f) and SYP (k-p) in the hippocampi of the NC and AD APOE ε4 noncarriers and carriers. Zoomed-in images of the CA1 and the entorhinal cortex (EC). (g-j, q-t) Representative images of immunohistochemical staining for SV2A (g-j) and SYP (q-t) in the frontal cortex (FC) and temporal cortex (TC) of AD patients. zoom-in images of gray matter (GM) and white matter (WM) in the cortex regions. Scale bars = 50 microns (b, d, f, l, n, p), 100 microns (j, t), and 400 microns (h, r).

**Fig. 4.**
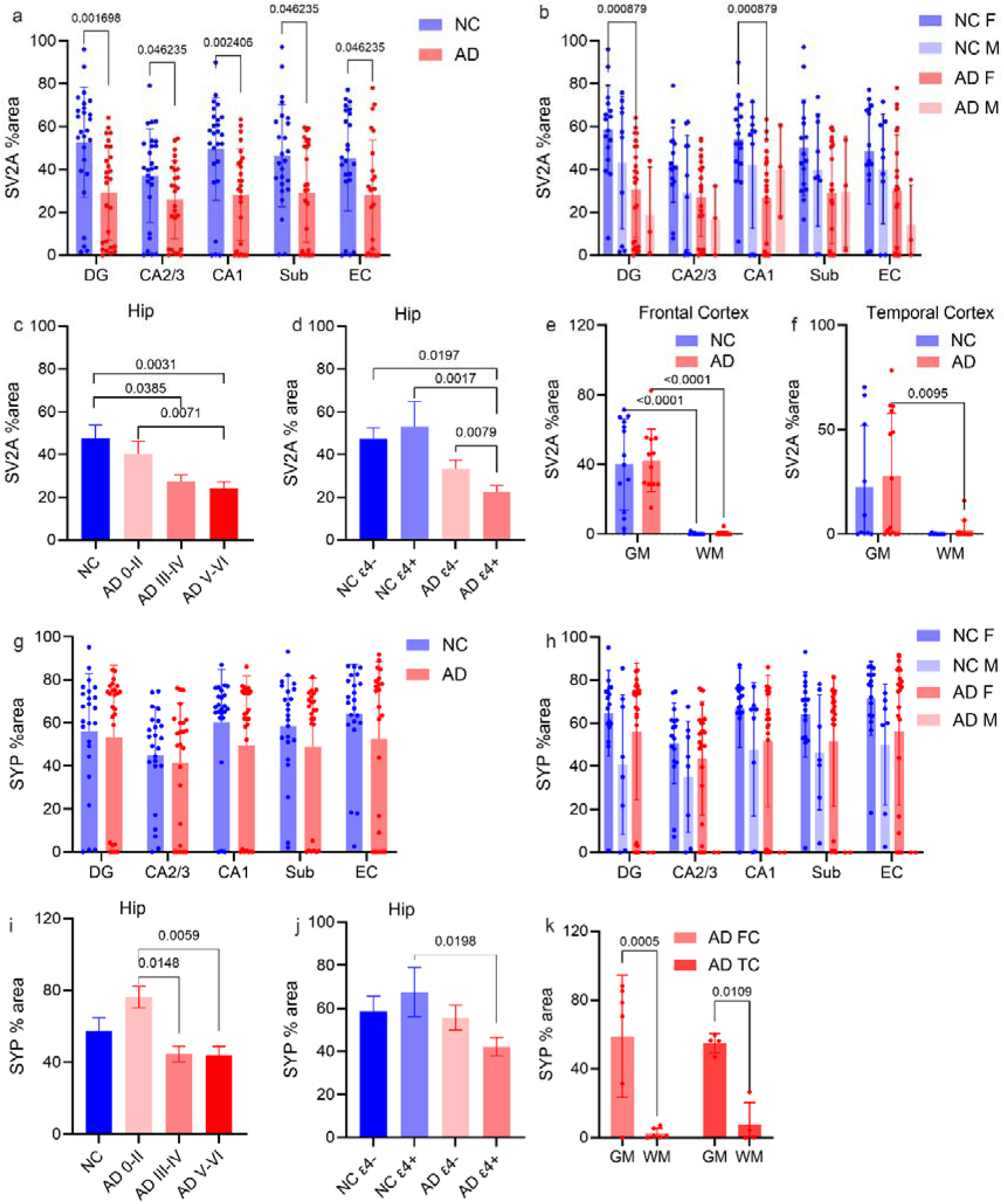
Reduced SV2A but not SYP levels in the hippocampus and entorhinal cortex of AD patients compared with NCs and associations with the APOE ε4 and Braak stages. (a, g) SV2A and SYP levels in the dentate gyrus (DG), CA3/CA2, CA1, and subiculum (Sub) of the hippocampus and the entorhinal cortex (EC) in AD patients compared with those in NCs and (b, h) in female and male AD patients and NCs. (c, i) Comparison of hippocampal SV2A and SYP levels between NC and AD patients at different Braak stages. (d, j) SV2A and SYP levels in APOE ε4 carriers and noncarriers of AD and NCs. (e, f, k) SV2A and SYP levels were greater in the gray matter (GM) than in the white matter (WM) of the frontal cortex (FC) or temporal cortex (TC).

The SYP level did not differ significantly between AD patients and NCs in the subfields of the hippocampus or the entorhinal cortex (**Fig. 3k-p**, **Fig. 4g-j**). Similar to SV2A, in the frontal cortex and temporal cortex of both AD patients and NCs, the level of SYP in gray matter was several-fold greater than that in white matter (**Fig. 3q-t**, **Fig. 4k**). Like SV2A, the laminar distribution of SYP in the gray matter of the frontal, temporal and entorhinal cortex is mostly homogenous; the pyramidal layer of the hippocampus shows dense SYP signals. For AD patients, NCs, and the combined cohort of AD patients and NCs, there was a strong positive correlation between SV2A and SYP in the hippocampus, entorhinal cortex and gray matter of the cortex (**Fig. 6j-p**).

Next, we assessed the influence of the APOE ε4 allele and sex differences in SV2A and SYP levels. We found that SV2A levels were lower in AD APOE ε4 carriers than in AD APOE ε4 noncarriers (**Fig. 3c-f**, **4d**, p=0.0079) but not in SYP carriers (**Fig. 3m-p**, **4j**). A lower level of SYP was observed in AD APOE ε4 carriers than in NC APOE ε4 carriers (**Fig. 4j**). There was no significant difference in the levels of SV2A or SYP between female and male AD patients or between female and male NCs (**Fig. 4c, h**).

### Regional levels of SV2A were positively correlated with SYP and negatively correlated with A**β**, phospho-tau, and the Braak stage

Greater levels of SV2A (**Fig. 4c**, p=0.0071) were detected in AD at Braak stages 0-II than in V-VI, as were greater levels of SYP in AD at Braak stages 0-II than in V-VI (**Fig. 4j,** p=0.0059) and III-IV (**Fig. 4j,** p=0.0148). Compared with those in the NCs, lower levels of SV2A were detected in the AD patients at Braak stages III-IV, as well as at Braak stages V-VI (**Fig. 4c,** p=0.0385, p=0.0031), but not in the AD patients at Braak stages 0--II. In contrast, no difference in the levels of SYP was detected between the NC and AD groups at different Braak stages (**Fig. 4j).**

Next, we explored the associations between the levels of Aβ and tau with SV2A and SYP in the hippocampus and entorhinal cortex of AD patients and NCs. 4G8 and AT8 staining was performed on the adjunct slices of the SV2A and SYP antibody-stained slices (**Fig. 5**). We found that there was no association between the CERAD-neuritic plaque score and the SV2A level or SYP level in the brains of AD patients and NCs. Negative correlations between SV2A and the 4G8+ Aβ level were detected in the DG (p=0.0122, r=-0.3748), CA3/CA2 (p=0.0384, r=-0.3246), and CA1 (p=0.0307, r=-0.3263) as well as in the entorhinal cortex (p=0.0051, r=-0.4448) (**Fig. 6a-d**) in the AD+NC groups. Robust negative correlations between the SV2A level and Braak stage were observed in the AD+NC group in the DG (p=0.0028, r=-0.9643), CA3/CA2 (p=0.0135, r=-0.8581), CA1 (p=0.0238, r=-0.8571), and Sub (p=0.0123, r=-0.8929) regions and in the entorhinal cortex (p=0.0238, r=-0.8571) (**Fig. 6e-i**, **STable 2**). In addition, SV2A and AT8 (phospho-Tau) levels were negatively correlated in the Sub (p=0.0326, r=-0.4374) of AD patients (**Fig. 6j, STable 2**). In contrast, no correlation was detected between SYP and 4G8 or AT8 levels or between SYP and the CERAD score or Braak stage in AD patients and NCs (**STable 2**).

**Fig. 5.**
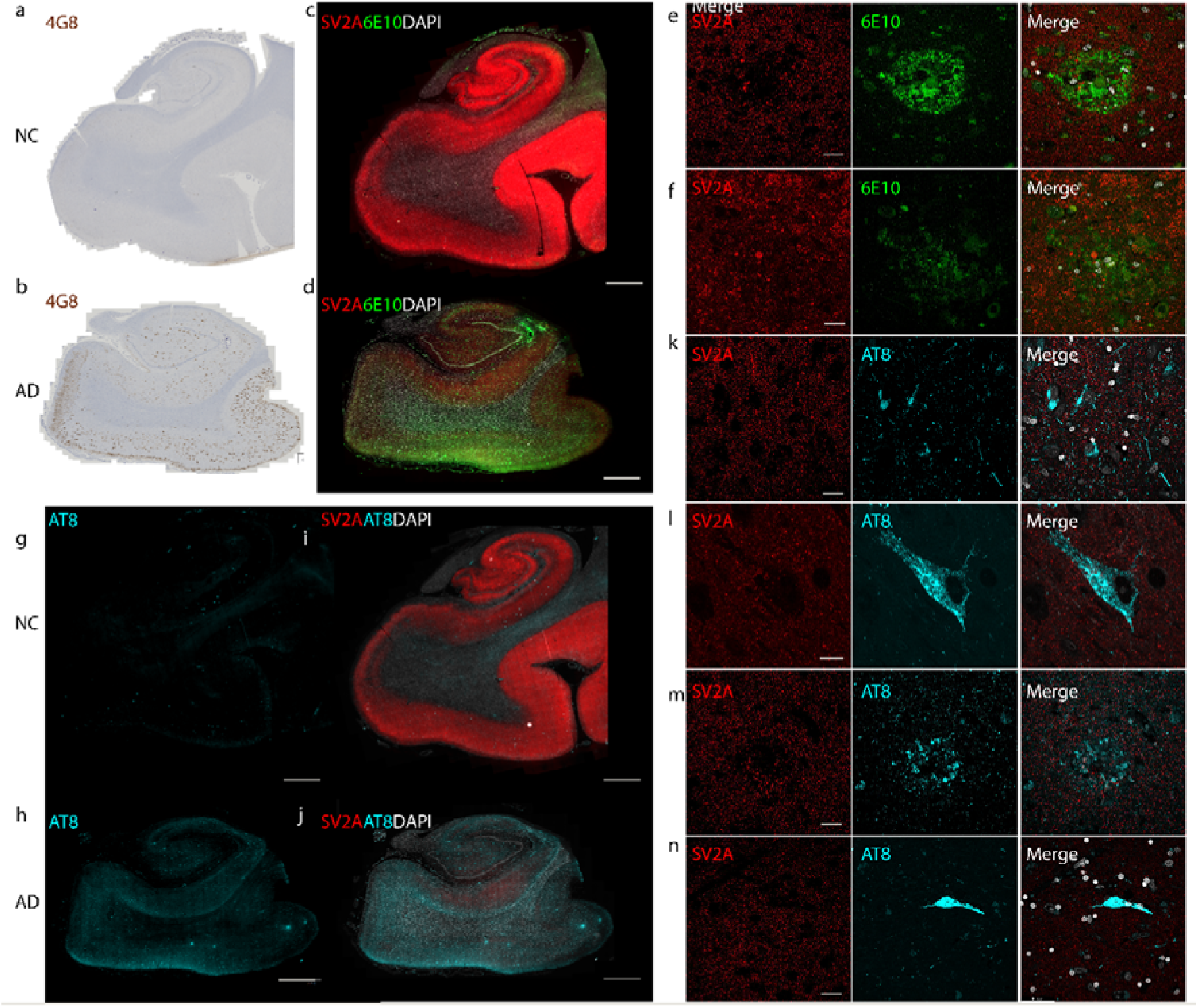
SV2A expression is lower in the presence of amyloid-β plaques (4G8, 6E10) and phospho-Tau (AT8) in AD patients than in NCs. Representative overview of 4G8 amyloid-β (brown) immunohistochemical staining in the NC and AD groups. (c-n) Representative overview and zoomed-in view of immunofluorescence staining of SV2A (red), 6E10 amyloid-β (green) and AT8 (cyan) in the hippocampi of the NC and AD groups. Nuclei were counterstained with DAPI (white). Core plaque (e), diffuse plaque (f), neuropil thread (n), mature tangle (l), neuritic plaque (m), and ghost tangle (n). Scale bars = 2 mm (c, d, g-j) and 20 microns (e, f, k-n).

**Fig. 6.**
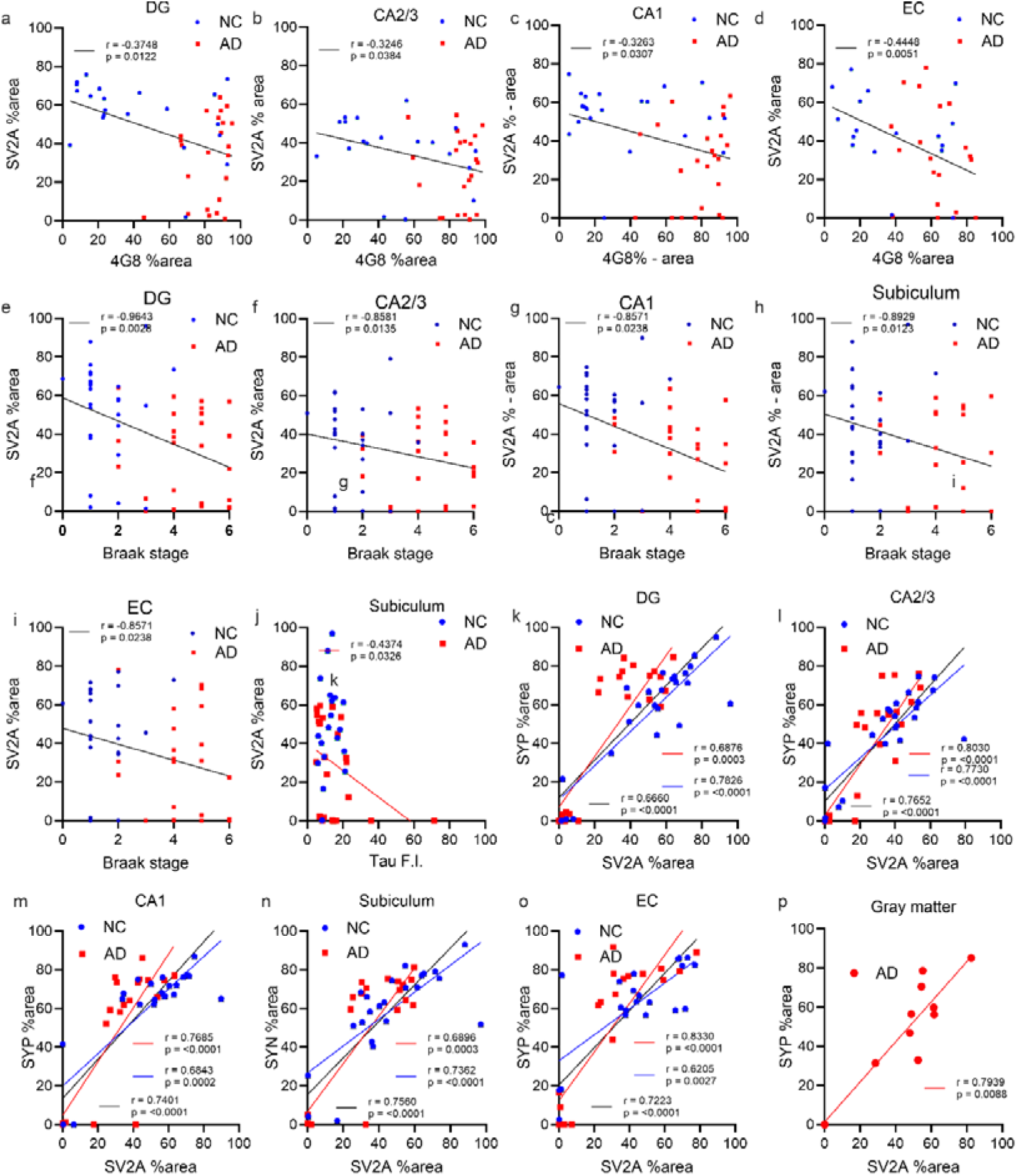
Correlation analysis between SV2A, amyloid-β, Braak stage and tau levels. (a-d) Negative correlations between 4G8 amyloid-β and SV2A levels in the hippocampus and EC in AD patients and NCs. (e-j) Negative correlations between AT8 (phospho-Tau), Braak stage and SV2A levels in the hippocampus and EC regions in AD+NCs. (k-p) Positive correlations between SYP and SV2A levels in different regions in AD patients and NCs. AD; red line, NC; black line, AD+NC. Detailed p and r values are provided in **STable 2**.

To further assess the associations between SV2A and different types/morphologies of Aβ and tau, double staining was performed for SV2A/6E10 and SV2A/AT8. A reduced level of SV2A was more evident in the cored plaque than in the diffuse plaque (**Fig. 5c-f**). Mature tangles, ghost tangles and neuritic plaques were associated with a reduced level of SV2A (**Fig. 5g-n**), whereas the presence of neuropil thread did not influence the intensity of SV2A staining. These results indicated a close association between SV2A with Aβ and tau pathologies.

## Discussion

One of the strengths of our study is the comprehensive analysis of relatively large samples of postmortem brain tissue from AD patients and NCs characterizing synaptic markers. We performed synaptosome proteomics in BdEVs from 17 AD patients and 4 NCs and immunostaining in 40 AD patients and 44 NCs. The observed alterations in synaptic proteins and their associations with Aβ, tau, the Braak stage and the APOE ε4 allele provide postmortem evidence for in vivo imaging and fluid biomarkers of synaptic density.

Here, we found reduced BdEV levels of SV2A and other synaptic proteins, such as SYT1, SNAP25, and 14-3-3 ζ, in AD patients compared with those in NCs and robust correlations among these synaptic markers. Interestingly, some of the markers, such as GAP43 and SYT1, were already reduced at the early Braak stage of AD, whereas others were reduced only at the late Braak stage of AD compared with those in the NC group, indicating that further analysis of fluid samples from a large cohort of patients at different disease stages is needed. An earlier study reported significantly lower levels of synaptic proteins in the plasma-isolated neural-derived exosomes of patients with AD than in those of NCs, which are correlated with cognitive decline (2). A recent study revealed altered CSF synaptic biomarkers such as 14-3-3 ζ/δ as promising pathophysiological changes in AD, whereas neuronal pentraxins are a general indicator of neurodegeneration and are associated with cognitive decline across various neurodegenerative dementias (53). In addition, we found negative correlations between SV2A and GFAP and the NEFL in the BdEV of AD patients. Synaptic biomarkers, such as neurogranin, NEFL, and SNAP25, have been shown to be positively associated with CSF and plasma levels of GFAP in AD patients (5, 85). A recent study revealed that the serum SV2A concentration was significantly negatively correlated with the serum GFAP concentration in a cognitively unimpaired population at risk of AD (80). [^18^F]SynVesT-1 measurements of brain SV2A levels were shown to be negatively associated with plasma GFAP and pTau181 in patients with cognitive impairment (83). In addition, CSF and serum levels of SV2A have been shown to be positively correlated with cognitive performance in patients with AD and decrease with the progression of AD (44, 80).

We demonstrated reduced levels of SV2A in the hippocampus and entorhinal cortex (but not in the frontal or temporal cortex) but no difference in the level of SYP between AD patients and NCs by using staining. The hippocampus is often the site with the earliest and most significant synaptic loss due to the degeneration of the entorhinal cortical cells that project to the hippocampus via the perforant path (7, 42, 59). Comparable protein expression levels of SV2A in the middle frontal gyrus in AD patients compared with NCs have been reported via ELISA and western blotting (47), whereas reduced levels of SV2A and SYP were reported by immunostaining (n=6–7 per group) in an earlier study (60). Different results have been reported via radioactive tracer binding assays and autoradiography, with lower (47) or unchanged (46) [^3^H]UCB-J levels in the middle frontal gyrus of AD patients than in NCs and unchanged [^3^H]UCB-J levels in the parietal, temporal, or occipital cortex of AD patients than in those of NCs (47). A recent immunoblotting study revealed reduced [^3^H]UCB-J but not SV2A protein levels in homogenates/different synaptosome fractions of the frontal cortex, temporal cortex and hippocampus of AD patients compared with those of NCs (32). In contrast, in vivo studies with different SV2A tracers revealed decreased uptake in the hippocampus and temporal cortex (3, 8, 9, 40, 43, 47, 73, 74), with divergent results (either no difference or reduction) in the frontal cortex in AD (and MCI) patients compared with NCs. These different observations from in vivo-ex vivo studies might be explained by the good correlation between SV2A autoradiography and PET but not between SV2A autoradiography and SV2A mRNA levels (26). In addition, in vivo SV2A uptake might be influenced by volumetric alterations as well as glucose metabolism (72), and the ex vivo SV2A level might be affected by postmortem delay.

The associations and distributions of SYP and SV2A are not fully understood. Here, we found a robust correlation between the quantification of SYP and SV2A expression by immunostainingof brain tissue slices from AD patients and NCs. This finding is in line with our measurement of the BdEV of the positive correlation between SYP and SV2A and with a recent publication in AD (60). In contrast, an earlier study in postmortem brain tissue from PD, PDD and DLB patients revealed that SYP and SV2A densities were both decreased but correlated only weakly despite their presynaptic localization in synaptic vesicles (20). One possible reason for this difference is the person-to-person variability in the synaptic density (50). In addition, we observed a generally homogenous pattern with a slight difference in the laminar distribution of SV2A density as well as in the SYP, similar to observations in earlier studies (20, 55, 60, 66).

APOE ε4 is the most important risk factor for sporadic AD and influences synaptic function according to studies in cell culture and animal models (34). Recent studies revealed that the APOE ε4 allele is associated with decreased SV2A PET uptake in the brains of cognitively impaired participants (21), in cognitively unimpaired APOE ε4/ε4 homozygotes (64), and with lower serum levels of SV2A (79, 80). Here, we provide postmortem evidence that AD APOE ε4 carriers presented lower SV2A levels in the hippocampus than did AD ε4 noncarriers. A study in E3FAD and E4FAD model mice showed that the detrimental effects of APOE ε4 might result from altered dendritic spine density and synaptic proteins (69). An earlier study showed that APOE ε4 markedly exacerbates tau-mediated neurodegeneration in a mouse model of tauopathy (63). APOE ε4 exerts a “toxic” gain of function, and the removal of astrocytic APOE decreases tau-induced synaptic loss and microglial phagocytosis of synaptic elements, suggesting a key role for astrocytic APOE in synaptic degeneration (76).

Moreover, we found negative associations between SV2A and p-Tau (AT8) and the Braak stage and between SV2A and Aβ (4G8) but not between the CERAD score in the hippocampus and entorhinal cortex of AD patients. This provides ground truth for in vivo imaging to evaluate the correlation between PET and fluid biomarker readouts for Aβ and between tau and SV2A. Earlier studies in disease animal models and in AD patients found that compared to Aβ. tau associated more closely with neurodegeneration and synaptic loss (4, 51, 81). Converging negative associations were observed in aMCI and AD patients in the hippocampus and entorhinal cortex via [^11^C]UCB-J with [^18^F]MK-6240 (74)/[^18^F]flortaucipir (45), [^18^F]SynVesT-1 with [^18^F]MK-6240 (37), and in the neocortex of AD patients via [^11^C]UCB-J with [^18^F]flortaucipir (12). Different results have been reported for Aβ deposits and SV2A from previous in vivo studies in AD patients: [^11^C]UCB-J revealed a "paradoxical" positive association between SV2A and local Aβ deposition ([^11^C]PiB) in the hippocampus of AD patients (54). Another PET study revealed a negative correlation between SV2A ([^18^F]SynVesT-1) and global Aβ deposition ([^18^F]florbetapir) in the bilateral hippocampus and parahippocampus of AD patients (37). In addition, our data suggest that the associations between SV2A and tau are much stronger than those between SV2A and Aβ. The accumulation of oligomeric tau and its spread from pre- to postsynapses has been found to be a cultiprite contributing to the spread of tau throughout the brain (10, 24). A postmortem study revealed a negative correlation between SV2A ([^3^H]UCB-J) and p-tau but not between SV2A and Aβ levels in the middle frontal gyrus in AD patients (46). Moreover, recent studies revealed that the serum and CSF levels of SV2A are negatively correlated with tau and p-tau but not with neurofilaments in cognitively unimpaired populations at risk of AD (14, 80), suggesting that distinct pathological pathways may be involved in synaptic and axonal degeneration.

There are several limitations in this study. First, there was an imbalance in sex in the AD group. Second, we did not stain using antibodies for Aβ oligomers, e.g., A11, or tau oligomers, e.g., T22, or other Tau antibodies, such as p-Tau217, which might provide additional insights. Third, the sample sizes used for BdEV proteomics were different from those used for staining and were smaller. Moreover, we did not divide the AD group into early-onset and late-onset AD groups.

## Conclusion

We demonstrated reduced levels of SV2A in the hippocampus and entorhinal cortex, as well as in BdEVs, in AD patients compared with NCs. The levels of SV2A positively correlate with other synaptic markers in BdEVs and with SYP in the brain and are negatively associated with Aβ, tau and the Braak stage. These results provide postmortem evidence that SV2A is a valuable marker for synaptic imaging and fluid biomarkers.

## Declaration

### Consent for participation

The consent for participation was collected by NBB and CHUV.

### Consent for publication

Consent for publication was obtained from NBB and CHUV.

## Acknowledgments

The authors acknowledge the NBB and CHUV Brain Banks for providing postmortem human brain tissue; the authors acknowledge the Center for Microscopy and Image Analysis (ZMB) for support with microscopy. The authors thank Serena Savodelli at ETH Zurich for their technical assistance and Dr. Manfredo Quadroni from the proteomics platform at the University of Lausanne for his advice and technical support.

## Author contributions

RN conceived and designed the study. JN performed the staining and confocal microscopy and quantified the microscopy data. AB, VZ, JE, LB, MC, and KR performed the proteomics and analysis. JN, KR, and RN wrote the first draft. All the authors contributed to the manuscript. All the authors have read and approved the final manuscript.

## Data availability

Data is available upon responable request from the corresponding authors.

