## Supplemental Files for "Reduced levels of synaptic vesicle protein 2A in the extracellular vesicles and brain of Alzheimer’s disease- associations with Aβ, tau and synaptophysin"

**STable 1. Antibodies and chemicals used for immunochemical staining**

| **Item** | **Catalog no** | **Dilution** | **Supplier** |
| --- | --- | --- | --- |
| Mouse phospho-Tau (Ser202, Thr205) monoclonal antibody (AT8) | MN1020 | 1:1000 | Invitrogen |
| Mouse purified anti-β-Amyloid, 17-24 monoclonal antibody (4G8, IHC) | 800701 | 1:4000 | Biolegend |
| Synaptophysin (27G12, IHC) | SYNAP-299-L-CE | 1:1000 | Leica Systems |
| Mouse purified anti-β-Amyloid, 1-16 monoclonal antibody (6E10, IF) | 803015 | 1:1000 | Biolegend |
| Anti-SV2A antibody (EPR23500-32, IHC, IF) | ab254351 | 1:1000 | Abcam |
| Alexa fluor488 donkey anti-mouse IgG (H+L) | 715-545-151 | 1:500 | Jackson |
| Alexa fluor674 donkey anti-rabbit IgG (H+L) | 711-605-152 | 1:500 | Jackson |
| Alexa fluor488 donkey anti-guinea pig IgG (H+L) | 706-545-148 | 1:500 | Jackson |
| Polymer Refine detection | DS9800 |  | Leica Systems |
| DAPI (4',6-Diamidino-2-Phenylindole, Dihydrochloride) | D1306 | 1:1000 | Invitrogen |

**STable 2. Correlation analysis of hippocampal SV2A and synaptophysin in AD patients and NCs**

|  | **Region** | **Group** | **SV2A** | | **SYN** | |
| --- | --- | --- | --- | --- | --- | --- |
|  |  |  | **r** | ***P* value** | **r** | ***P* value** |
| Braak Stage | CA1 | NC | 0.1000 | 0.9500 | 0.2000 | 0.9167 |
|  |  | AD | -0.3000 | 0.6833 | -0.4000 | 0.5167 |
|  |  | NC+AD | **-0.8571** | **0.0238** | -0.6000 | 0.2417 |
|  | CA2/3 | NC | -0.4919 | 0.3999 | 0.2000 | 0.9167 |
|  |  | AD | 0.0776 | 0.9100 | -0.4000 | 0.5167 |
|  |  | NC+AD | **-0.8581** | **0.0135** | -0.6571 | 0.1750 |
|  | DG | NC | 0.1000 | 0.9500 | 0.2000 | 0.9167 |
|  |  | AD | -0.4000 | 0.5167 | -0.4000 | 0.5167 |
|  |  | NC+AD | **-0.9643** | **0.0028** | -0.6000 | 0.2417 |
|  | SUB | NC | 0.6000 | 0.3500 | 0.2000 | 0.9167 |
|  |  | AD | -0.4000 | 0.5167 | -0.4000 | 0.5167 |
|  |  | NC+AD | **-0.8929** | **0.0123** | -0.6000 | 0.2417 |
|  | EC | NC | 0.4000 | 0.5167 | 1.0000 | 0.0833 |
|  |  | AD | -0.3000 | 0.6833 | -0.4000 | 0.5167 |
|  |  | NC+AD | **-0.8571** | **0.0238** | -0.7714 | :0.1028 |
| Tau (AT8) | CA1 | NC | 0.09474 | 0.6912 | -0.06708 | 0.7914 |
|  |  | AD | -0.1992 | 0.3397 | -0.2709 | 0.1903 |
|  |  | NC+AD | -0.02108 | 0.8907 | -0.2058 | 0.1854 |
|  | CA2/3 | NC | -0.04361 | 0.8551 | 0.1146 | 0.6508 |
|  |  | AD | -0.2654 | 0.1998 | -0.2577 | 0.2135 |
|  |  | NC+AD | -0.1337 | 0.3812 | -0.1549 | 0.3214 |
|  | DG | NC | 0.01053 | 0.9649 | -0.003096 | 0.9903 |
|  |  | AD | -0.2260 | 0.2670 | -0.01880 | 0.9274 |
|  |  | NC+AD | 0.09911 | 0.5123 | -0.05722 | 0.7122 |
|  | SUB | NC | 0.1386 | 0.5715 | -0.004902 | 0.9887 |
|  |  | AD | **-0.4374** | **0.0326** | -0.3122 | 0.1375 |
|  |  | NC+AD | -0.1808 | 0.2460 | -0.2016 | 0.2063 |
|  | EC | NC | **0.5539** | **0.0230** | 0.3250 | 0.2370 |
|  |  | AD | -0.1887 | 0.3884 | -0.02471 | 0.9109 |
|  |  | NC+AD | 0.1240 | 0.4458 | 0.06741 | 0.6876 |
| Aβ (4G8) | CA1 | NC | -0.2208 | 0.3362 | -0.1632 | 0.5045 |
|  |  | AD | 0.2905 | 0.1787 | 0.02429 | 0.9146 |
|  |  | NC+AD | **-0.3263** | **0.0307** | -0.2187 | 0.1696 |
|  | CA2/3 | NC | -0.3877 | 0.1010 | -0.3235 | 0.2050 |
|  |  | AD | 0.01186 | 0.9682 | -0.04677 | 0.8405 |
|  |  | NC+AD | **-0.3246** | **0.0384** | -0.1740 | 0.2962 |
|  | DG | NC | -0.3909 | 0.0797 | -0.08246 | 0.7372 |
|  |  | AD | 0.2421 | 0.2657 | -0.1779 | 0.4168 |
|  |  | NC+AD | **-0.3748** | **0.0122** | -0.6950 | 0.6950 |
|  | SUB | NC | -0.2501 | 0.2615 | -0.1489 | 0.5310 |
|  |  | AD | 0.09289 | 0.6734 | 0.06028 | 0.7847 |
|  |  | NC+AD | -0.2362 | 0.1182 | -0.1265 | 0.4187 |
|  | EC | NC | -0.3627 | 0.1529 | -0.2647 | 0.3207 |
|  |  | AD | -0.3988 | 0.0733 | :-0.2998 | 0.1990 |
|  |  | NC+AD | **-0.4448** | **0.0051** | -0.2934 | 0.0824 |
| SYN | CA1 | NC | **0.6843** | **0.0002** |  | |
|  |  | AD | **0.7685** | **<0.0001** |  |  |
|  |  | NC+AD | **0.7401** | **<0.0001** |  |  |
|  | CA2/3 | NC | **0.7730** | **<0.0001** |  |  |
|  |  | AD | **0.8030** | **<0.0001** |  |  |
|  |  | NC+AD | **0.7652** | **<0.0001** |  |  |
|  | DG | NC | **0.7826** | **<0.0001** |  |  |
|  |  | AD | **0.6876** | **0.0003** |  |  |
|  |  | NC+AD | **0.6660** | **<0.0001** |  |  |
|  | SUB | NC | **0.7362** | **<0.0001** |  |  |
|  |  | AD | **0.6896** | **0.0003** |  |  |
|  |  | NC+AD | **0.7560** | **<0.0001** |  |  |
|  | EC | NC | **0.6205** | **0.0027** |  |  |
|  |  | AD | **0.8330** | **<0.0001** |  |  |
|  |  | NC+AD | **0.7223** | **<0.0001** |  |  |

Analysis of immunohistochemical staining of in the hippocampus of 40 AD, 44 NC; AD, Alzheimer’s disease. NC, nondemented control group. Nonparametric Spearman’s rank correlation was used. r: Correlation coefficient. SYN, synaptophysin. No correlation between CERAD score with SV2A or synaptophysin was detected.

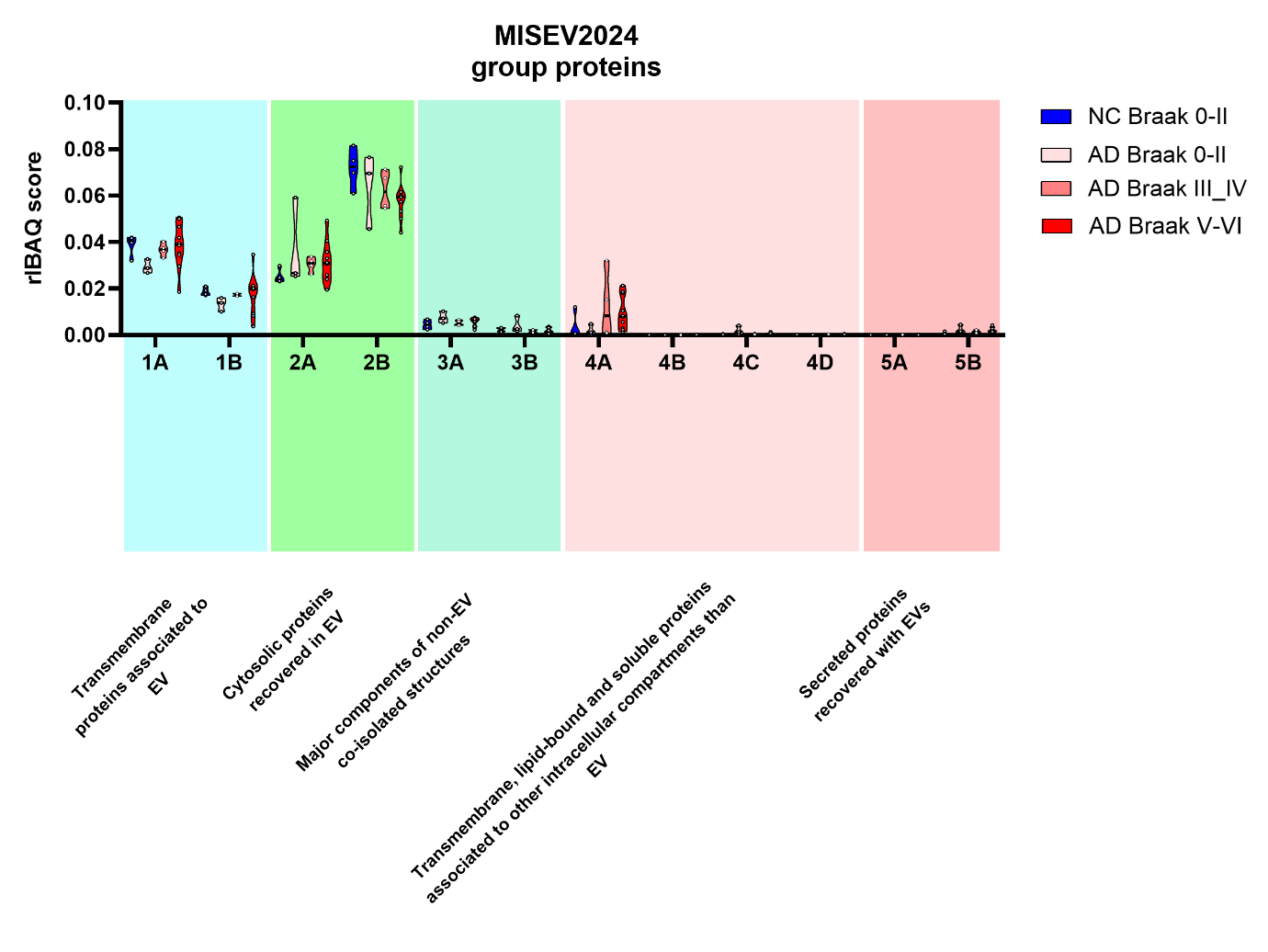

**SFig. 1 MISEV group proteins in NC and AD BdEVs.**

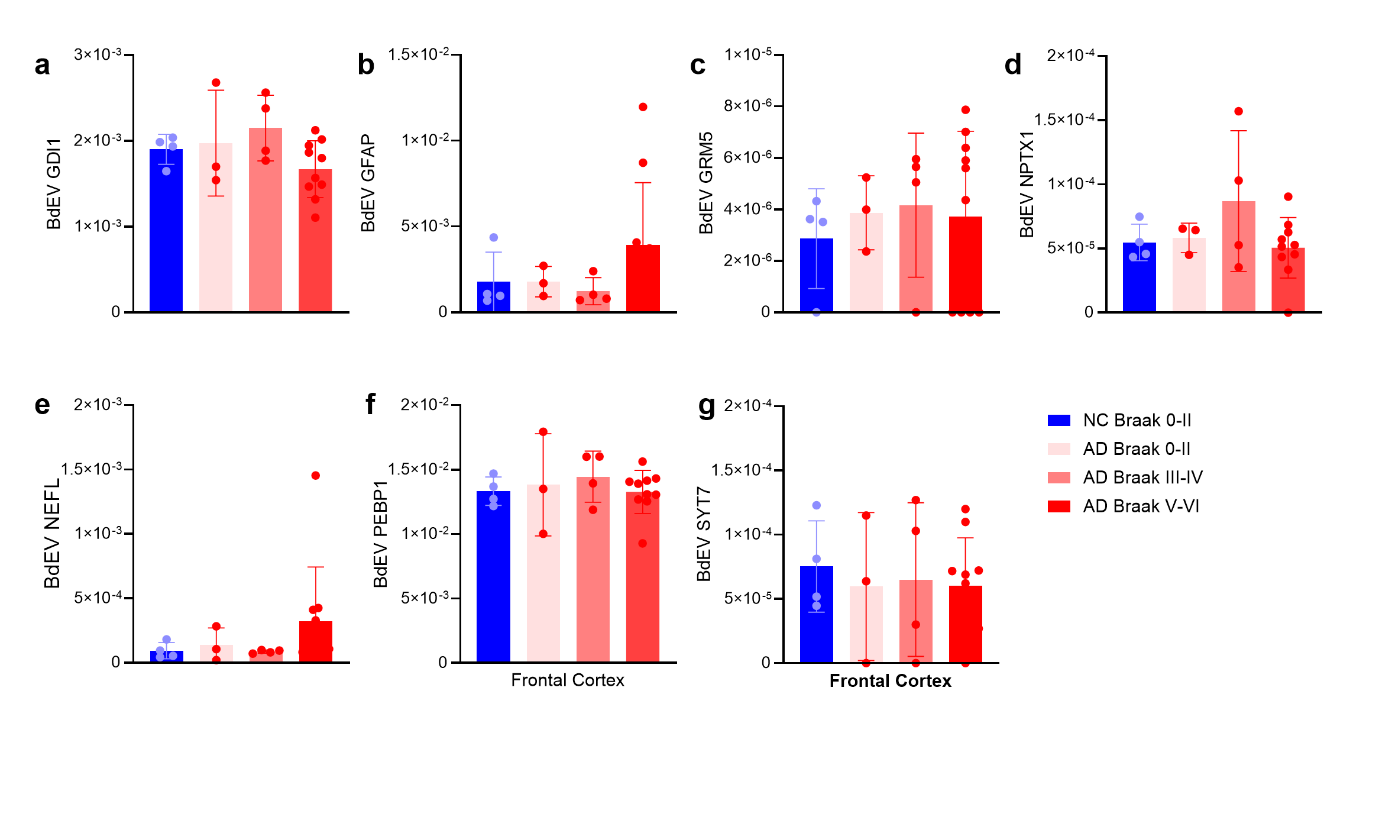

**SFig. 2 Comparison and correlation between cortical synaptosome BdEV markers in NC and AD** (a-g) Comparison of BdEV levels of GDI1, GFAP, GRM5, NPTX1, NEFL, PEBP1, and SYT7 in the NC Braak 0-II, AD of Braak 0-II, III-IV, and V-VI.

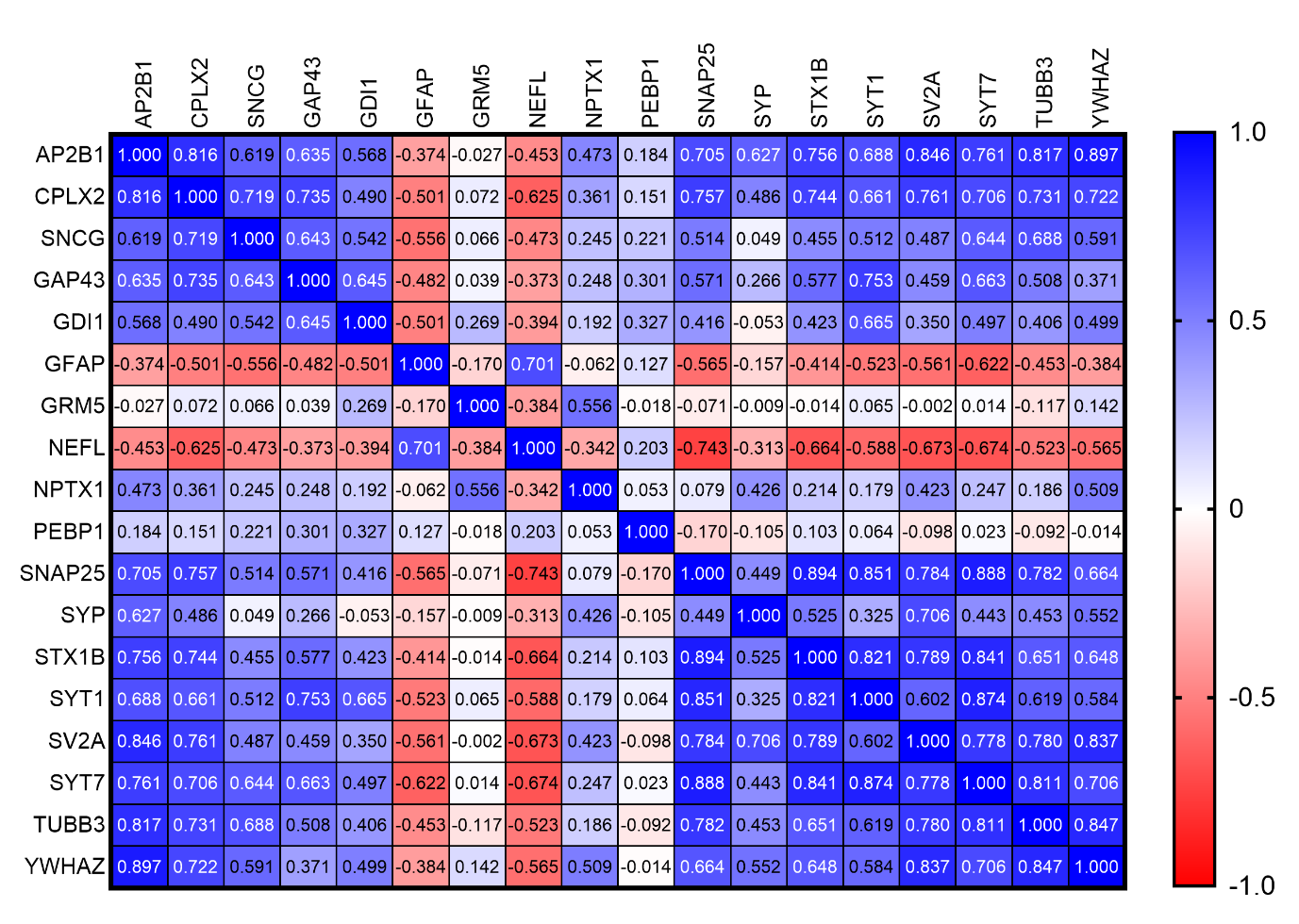

**SFig. 3 Nonparametric Spearman rank analysis of the rIBAQ matrix of correlations in the AD and NC** groups. The value in each cell indicates the p value.

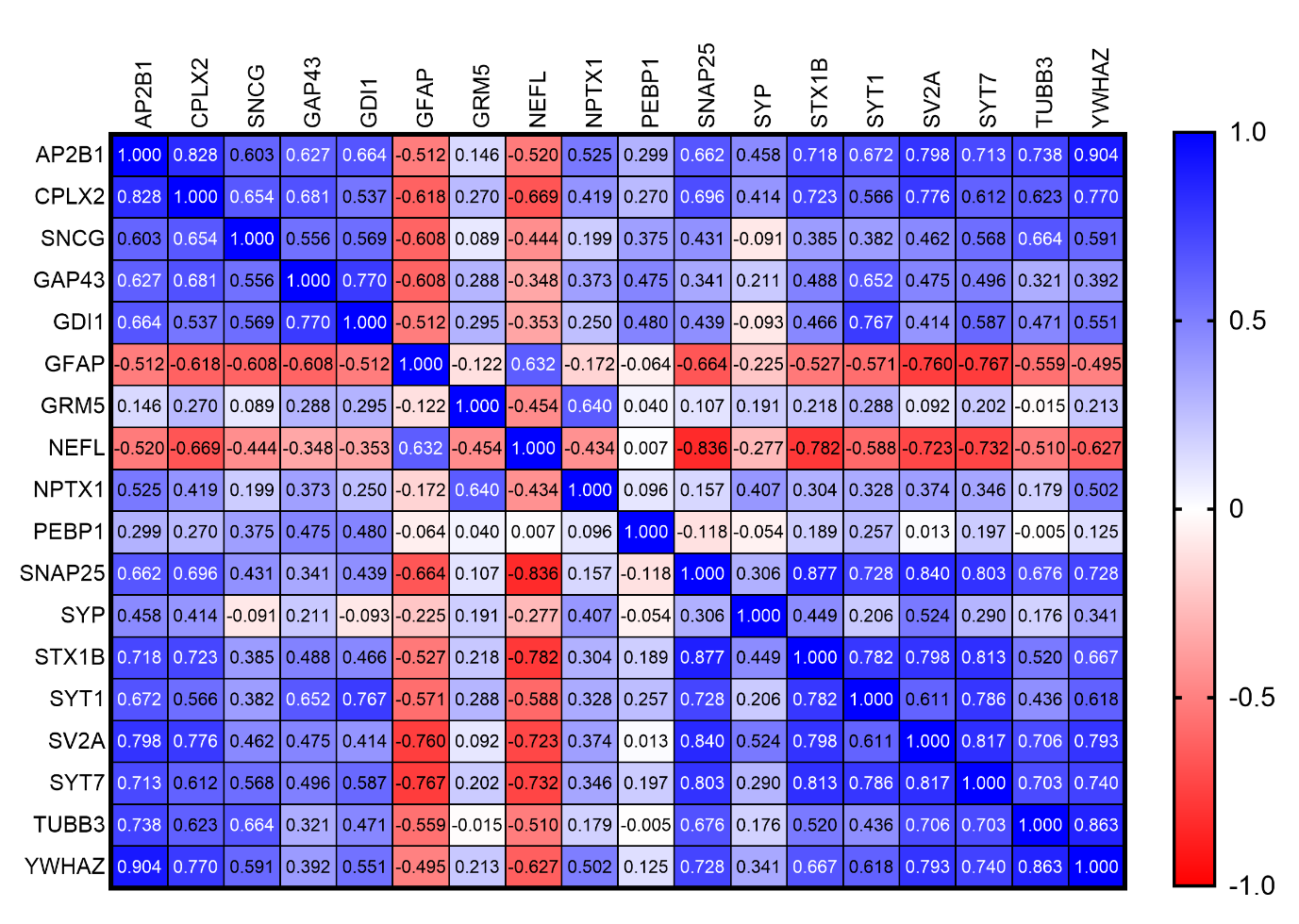
**SFig. 4 Nonparametric Spearman rank analysis of the rIBAQ matrix of correlation in the AD group**. The values in each cell indicate the p value

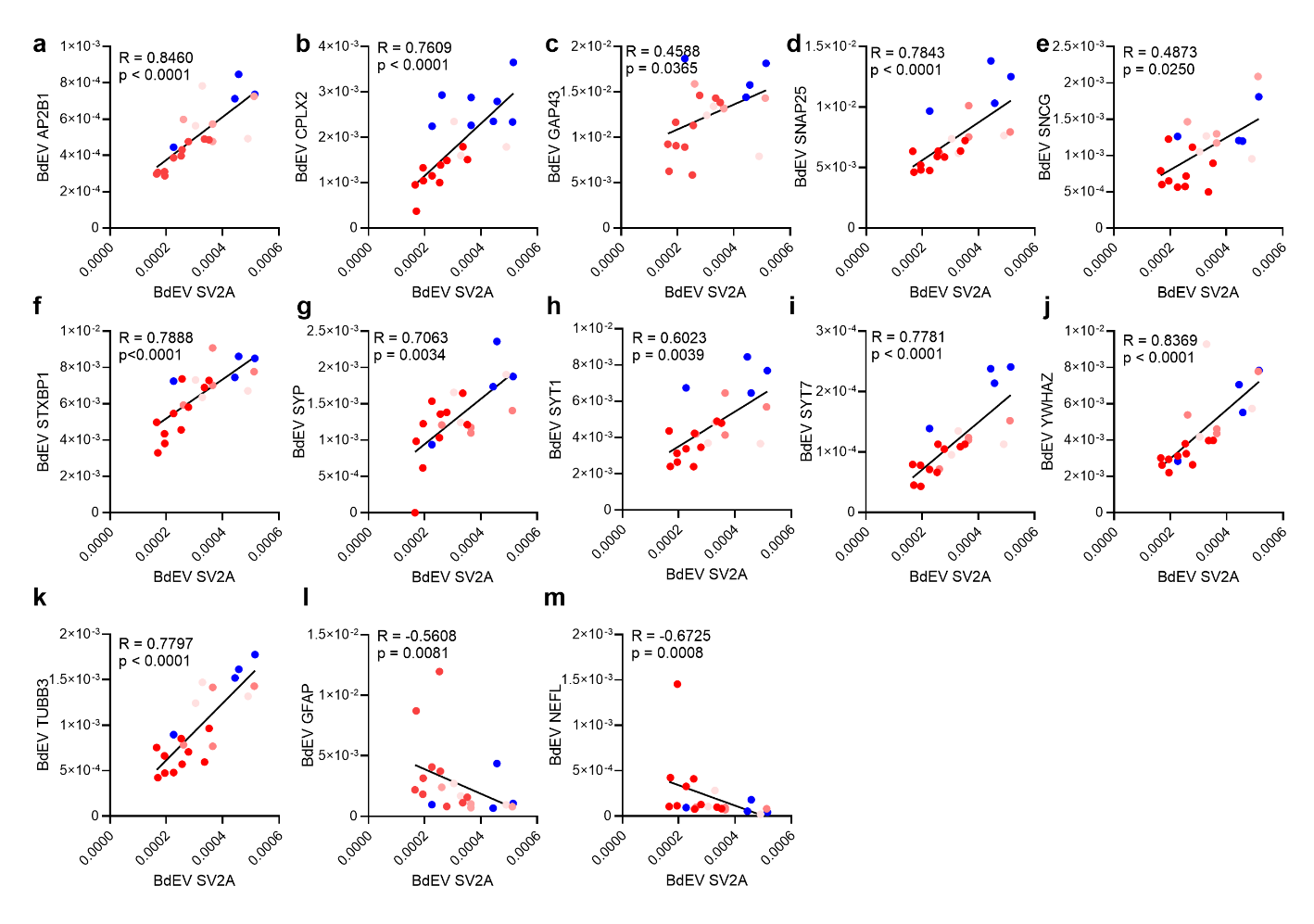

**SFig. 5 Nonparametric Spearman rank analysis of SV2A with other BdEVs in the AD and NC groups**

AP2B1, CPLX2, GAP43, SNAP25, SNCG, STX1B, SYP, SYT1, SVT7, YWHAZ, TUBB3, GFAP, and NEFL.

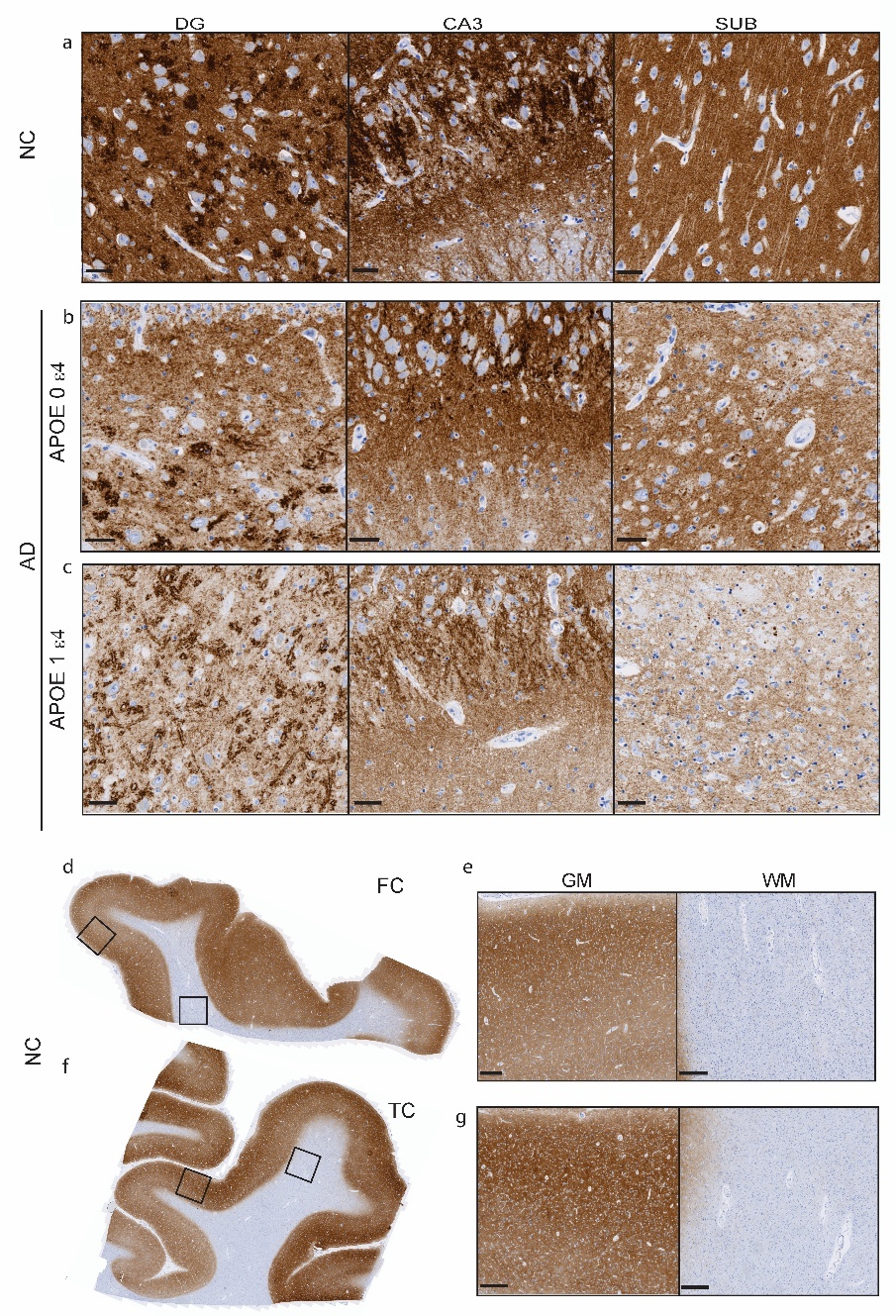

**SFig. 6 SV2A staining in the hippocampus, frontal cortex and temporal cortex of NC and AD APOE ε4 carriers and noncarriers.** (a-c) Zoomed-in images of immunohistochemical staining for SV2A (a-f) in the hippocampi of NC and AD APOE ε4 noncarriers and carriers. (d-g) Representative images of immunohistochemical staining for SV2A (g-j) in the frontal cortex (FC) and temporal cortex (TC) of the NC group. zoom-in images of gray matter (GM) and white matter (WM). Scale bars = 50 microns (a-c) and 400 microns (e, g).

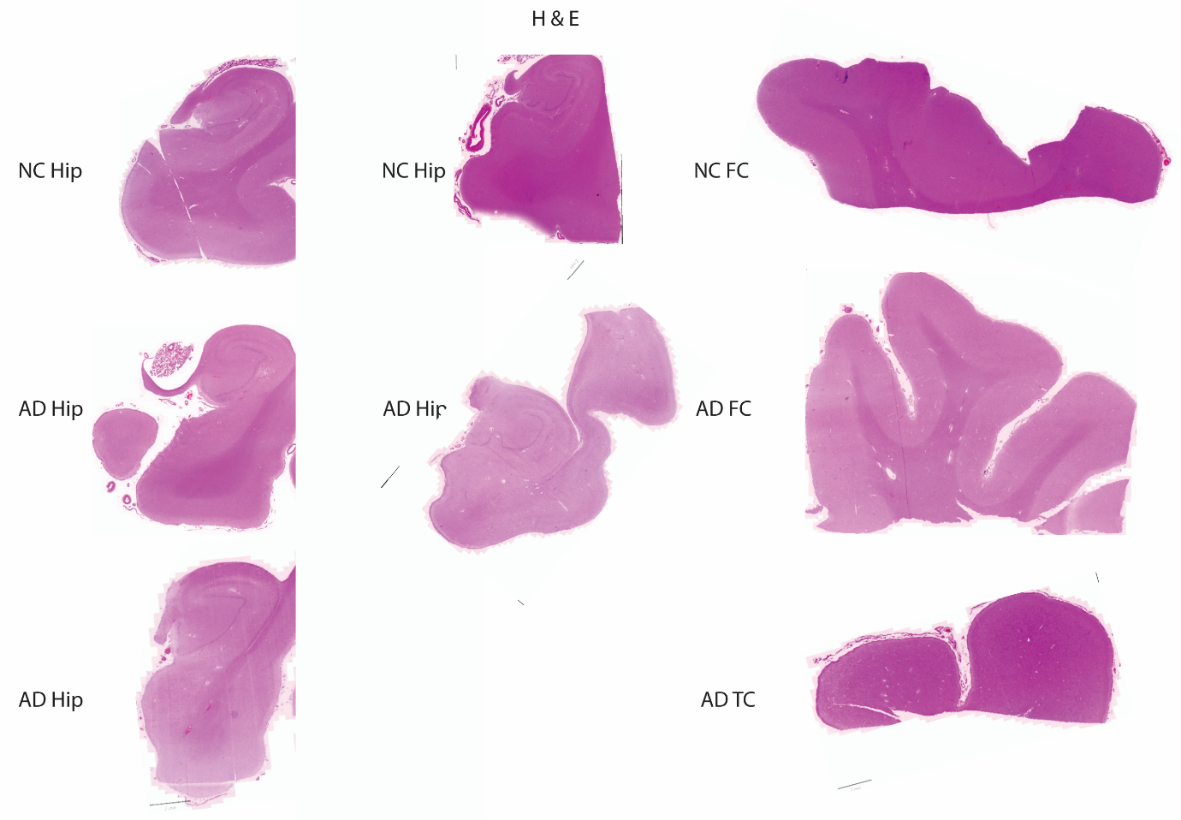

**SFig. 7 Representative images of H&E-stained images of the hippocampus, entorhinal cortex, frontal cortex and temporal cortex of AD patients and NCs.** Hip: hippocampus; H&E, hematoxylin and eosin; FC: frontal cortex; TC: temporal cortex.

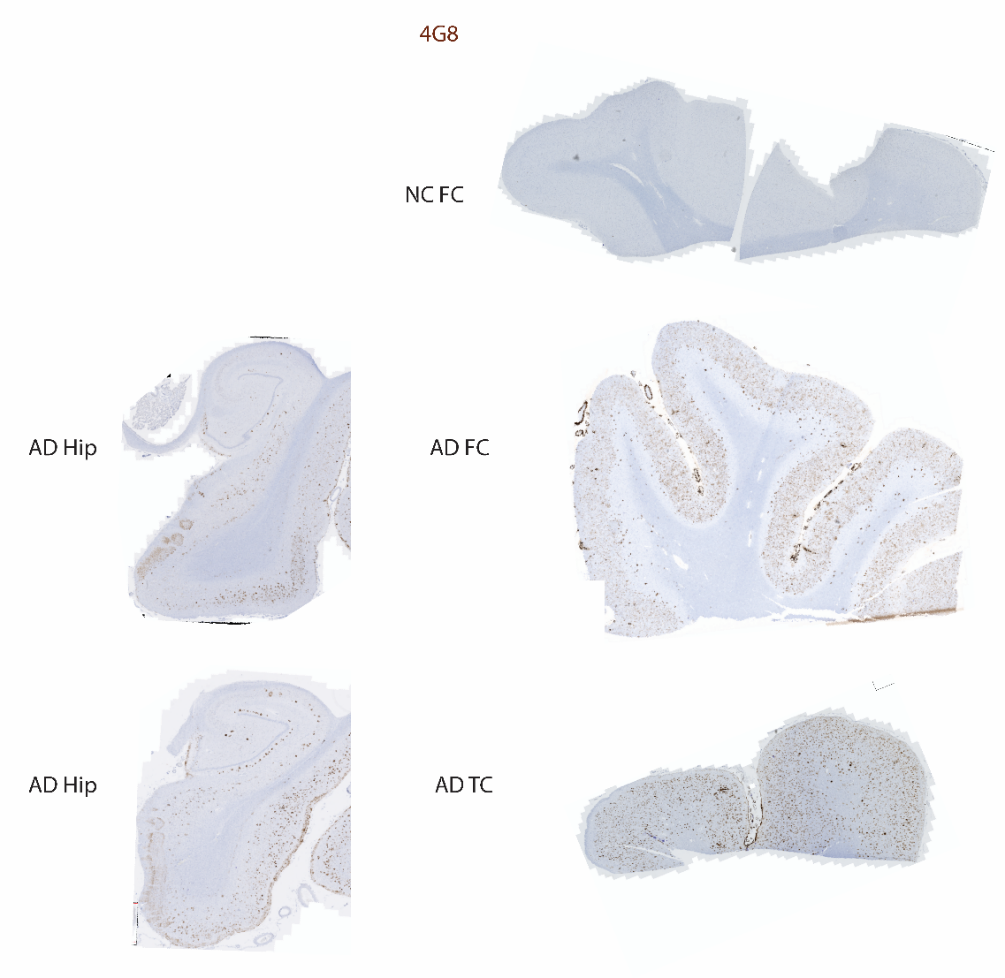

**SFig. 8** **Representative overview of 4G8 amyloid-β (brown) immunohistochemical staining in the NC and AD groups.**
